# Lung-resident memory B cells maintain allergic IgE responses in the respiratory tract

**DOI:** 10.1101/2024.02.20.581262

**Authors:** Alexander J. Nelson, Bruna K. Tatematsu, Jordan R. Beach, Dorothy K. Sojka, Yee Ling Wu

## Abstract

Allergen-specific IgE is a key mediator of allergic asthma. However, the tissue sites and cell types that support IgE production at the mucosa remained undefined. Here, we reveal that inhaled allergens induce the formation of IgG1^+^ lung-resident memory B cells (MBC) that switch to IgE. Using a mouse reporter for IgE class switch recombination, a requirement for the generation of IgE, we identify lung tissues as a major site of IgE class switching, which is dominated by IgG1^+^ MBCs and supported by IL-4-producing T_H_2 cells. This is in sharp contrast to what occurs in the draining lymph nodes where germinal center B cells are the IgE-switching population and T_FH_ cells provide IL-4. By single-cell transcriptomic analyses, we reveal potential mechanisms for the formation of lung-resident MBCs. Altogether, this study identifies the origin of allergen-specific IgE in the respiratory tract and explains how local chronic hypersensitivity is maintained in allergic asthma.

## INTRODUCTION

Immunoglobulin E (IgE) is a potent mediator of allergic diseases including allergic asthma^1,2^. Sensitization to allergens in the airway drives the generation of allergen-specific IgE that mediates the degranulation of effector cells such as mast cells, leading to the development of airway hypersensitivity and pathology. Such hypersensitivity is maintained by immunological memory and lasts for years, with recurrent exposure to allergens driving the intermittent nature of allergic asthma symptoms.

Immunological memory for IgE is distinctive from other antibody isotypes. Typically, antigen-specific antibodies are maintained by long-lived plasma cells and memory B cells (MBCs). Plasma cells in the bone marrow maintain systemic antibody titers and MBCs circulate in the blood and localize to lymphoid organs^3–6^. Memory B cells are quiescent but can rapidly initiate robust secondary immune responses upon re-encounter of antigens by seeding the formation of new germinal centers and differentiating into antibody-secreting plasmablasts^5,7–9^. However, IgE-expressing B cells minimally contribute to the memory compartment^10^. Long-lived IgE-producing plasma cells are rare and typically only observed in humans during severe allergies or persistent allergen exposure^11^ and in experimental models in which animals were systemically immunized or subjected to frequent allergen challenges for several weeks^12,13^. Furthermore, IgE-expressing B cells are transient, prone to apoptosis, and fail to form MBCs^14–16^, raising the question of how allergen-specific IgE responses are maintained long-term.

Studies in humans showed that some locally allergic patients display IgE in the respiratory tract even in the absence of detectable systemic IgE^17,18^, suggesting systemic IgE is not required for local allergic responses but rather IgE memory may be maintained locally in the sensitized respiratory tract. This is further supported by analyses of bronchial biopsies from allergic asthma patients revealing the presence of allergen-specific IgE and ε heavy chain transcripts, providing evidence for the presence of IgE secreting B cells in the sensitized mucosa^19,20^. However, it was unclear whether IgE-producing B cells are generated in lymph nodes and travel to the respiratory tract, or whether exposure to allergens induces the formation of IgE-producing B cells by a long-lived cellular precursor in the respiratory mucosa. Since IgE^+^ MBCs do not exist, antigen-experienced, non-IgE expressing MBCs capable of undergoing class switch recombination (CSR) have been implicated as the potential cellular source of pathogenic IgE in airway hypersensitivity^19,21–24^. Indeed, switch circles, or DNA excised from the immunoglobulin heavy chain loci during CSR in the form of circles, were detected in lung biopsies from both atopic and non-atopic asthma patients^23^, indicating active IgE CSR at the lung mucosa. The origins of these MBCs and the mechanisms that promote CSR in allergen sensitized lungs remain to be carefully characterized.

Naïve B and T lymphocytes circulate through lymph and blood. During infection, naïve lymphocytes with specificity for antigens from the pathogen interact and become activated in secondary lymphoid tissues – lymph nodes and the spleen. Some of these antigen-experienced lymphocytes develop into memory cells, which continue to patrol through the circulation and tissues. Some of these lymphocytes, however, can be retained as “tissue-resident” memory cells in peripheral tissues such as the respiratory mucosa where re-encounter with the same microbes is likely. Tissue-resident CD8^+^ and CD4^+^ memory T cells capable of mounting rapid effector functions locally are well described in viral and bacterial infections^25^. In the past few years, tissue-resident MBCs have been reported in the lung after infection with influenza or *Streptococcus pneumoniae*^26,27^. These tissue-resident MBCs are the primary source of neutralizing antibodies in the airway, protecting the host from re-infection^26–28^. It is currently unknown whether tissue-resident MBCs are unique to local immune responses in T_H_1-biased responses against pathogens, or whether repeated allergen exposure in T_H_2-biased inflammation such as allergic asthma may similarly lead to the formation of tissue-resident MBCs and account for the persistence of hypersensitivity in the respiratory tract.

In this study, we show that B cells accumulate in the lungs after allergen inhalation and that MBCs persist as resident cells. Using fluorescent reporter mice for IgE switching, we identified lung-resident allergen-specific IgG1^+^ MBCs as the major source of IgE switching cells. These MBCs expand after antigen rechallenge and mediate local IgE responses in the airway. Our study demonstrates IgE memory is maintained by lung-resident MBCs at the sensitized respiratory mucosa.

## RESULTS

### Local allergen exposure elicits IgE responses in the lungs

In susceptible individuals, exposure to allergens in the airway results in the development of allergic asthma^29^. To investigate B cell responses to inhaled allergens, we used repeated intranasal administration of ragweed pollen (RWP), a relevant human respiratory allergen^30^, and ovalbumin (OVA), which enables the detection of antigen-specific antibodies and B cells **(Figure 1A)**. We administered RWP and OVA for 5 consecutive days in the first week to initiate allergic inflammation, then OVA alone for two weeks to boost cellular and humoral responses to OVA. As a control, we administered PBS alone intranasally for 3 weeks. In mice receiving RWP+OVA, but not PBS control mice, we observed hallmarks of allergic airway inflammation: infiltration of eosinophils into the airway and lung (**Figures 1B and S1A**) and induction of local antigen-specific IgE and IgG1 in the bronchoalveolar lavage fluid (BALF) (**Figure 1C**), as well as circulating IgE and IgG1 in serum (**Figure S1B**). Profiling of B cell responses to allergen exposure by flow cytometry revealed the emergence of germinal center (GC) B cells and T follicular helper (T_FH_) cells in lung-draining mediastinal lymph nodes (medLN), but no increase in GC B cells or T_FH_ cells in the spleen, indicating a localized response to allergen (**Figure S2**). Together with the presence of OVA-specific IgE in BALF, we hypothesized that local IgE responses in the airway may be mediated by B cells that infiltrate into the lungs.

**Figure 1.**
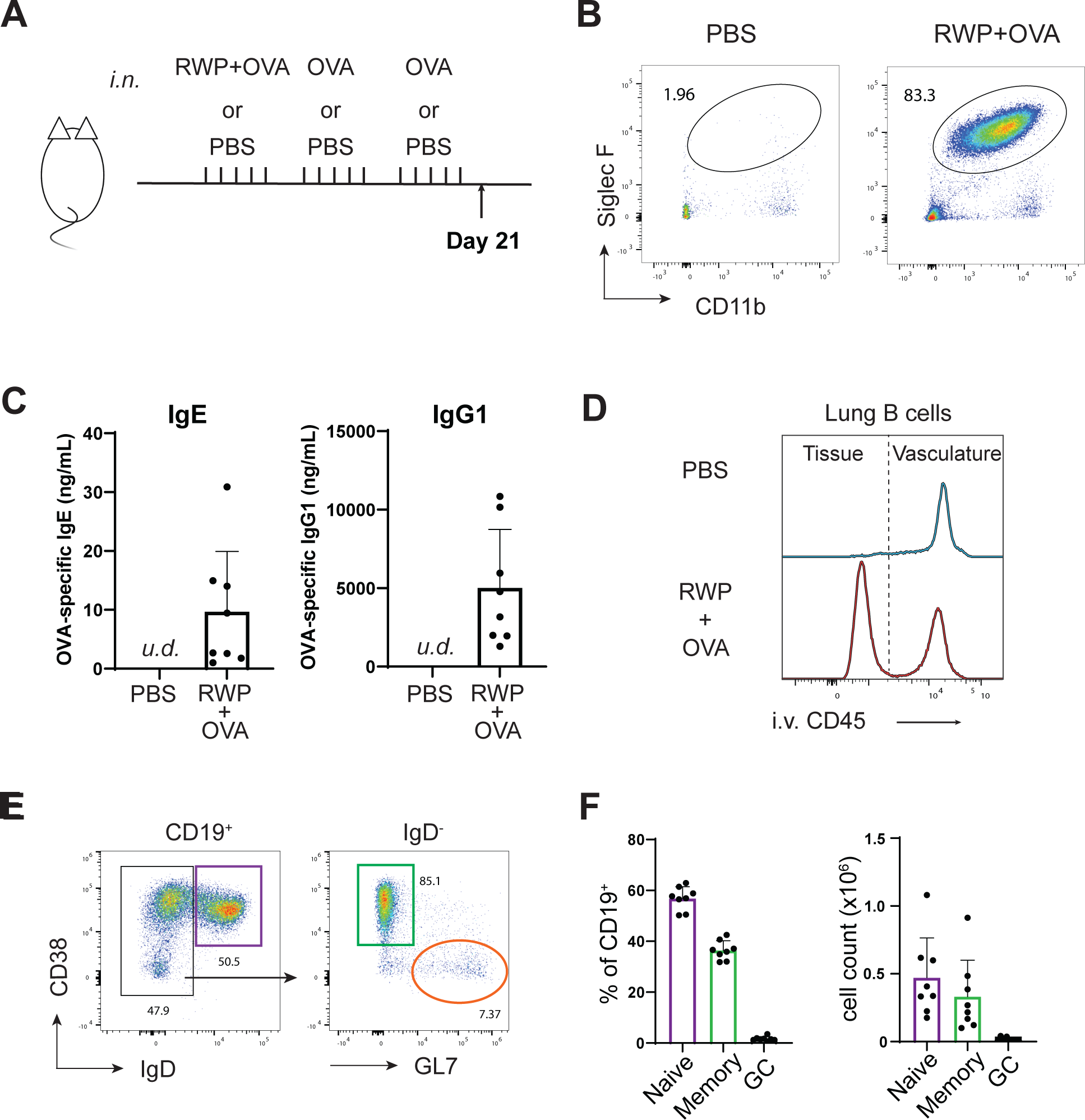
Mouse model of allergic airway inflammation. **(A)** Schematic of local allergen sensitization. Mice were treated intranasally with ragweed pollen and ovalbumin (RWP+OVA) for five days in the first week, followed by two weeks of OVA alone, and vehicle control mice received intranasal PBS. Mice were analyzed on day 21. **(B)** Eosinophils (live CD45^+^ CD11c^-^ CD11b^+^ SiglecF^+^) in bronchoalveolar lavage fluid (BALF) of PBS control (left) and allergen-sensitized (RWP+OVA) mice. Numbers are frequency, representative of 3 independent experiments with 3-8 mice per experiment. **(C)** Measurement of OVA-specific IgE and IgG1 in BALF by ELISA. **(D)** Intravascular labeling of lung B cells (live B220^+^) with anti-CD45 showing tissue-localized (unlabeled) and blood B cells (labeled). **(E)** Representative flow cytometry plots showing gating strategy for the identification of naïve (CD38^+^ IgD^+^, purple), memory (IgD^-^ CD38^+^ GL7^-^, green), and germinal center (GC, IgD^-^ CD38^-^ GL7^+^, orange) B cells (live B220^+^ i.v. CD45^-^) in the lungs of allergen-sensitized mice. **(F)** Frequencies and cell counts of naïve, memory, and GC B cells in the lungs. Representative data from four independent experiments with 3-8 mice per group in each experiment (mean ± SD).

To identify lung-localized B cells by flow cytometry, we utilized intravenous injection of fluorophore-labeled anti-CD45 prior to sacrifice to distinguish B cells in the lung blood vessels (labeled, i.v. CD45^+^) from those in the lung tissue (i.v. CD45^-^) and therefore protected from fluorophore labeling. In mice receiving RWP+OVA, but not in PBS control mice, we observed lung-localized B cells protected from labeling (**Figure 1D**), indicating that allergen inhalation induces B cell infiltration into the lungs. Lung-localized B cells were mostly naïve (CD38^+^GL7^-^IgD^+^, 57 ± 4%) or memory phenotype (CD38^+^GL7^-^IgD^-^, 36 ± 3%) with a smaller population of GC phenotype B cells (CD38^-^IgD^-^GL7^+^, 1.6 ± 0.8%) (**Figures 1E and 1F**). Together, these data reveal that repeated allergen inhalation induces a local immune response involving infiltration of B cells into the lungs and allergen-specific IgE in the airway.

### Memory B cells are major IgE-switching cells in allergic lungs

IgE is produced when activated B cells undergo class switch recombination (CSR) to IgE. Due to the rare nature of these events, the tissue sites that support IgE CSR are poorly understood. To this end, we measured IgE CSR using Iε-tdTomato mice, which report germline transcription at the Iε locus, a prerequisite and molecular marker for IgE CSR, with the expression of the fluorescent protein tdTomato^31^ (**Figure 2A**). After three weeks of allergen inhalation, we profiled the expression of tdTomato in B cell subsets in the lungs, lung-draining mediastinal lymph nodes (medLNs), and spleen. Iε-tdTomato^+^ B cells were detectable in each tissue, with the highest frequency in the lungs and medLNs, and the lowest in the spleen (**Figure 2B**). Despite their low frequency, the number of Iε-tdTomato^+^ B cells was the highest in the spleen due to the relatively large number of B cells in the spleen compared to the lungs or medLN (**Figure 2B**). Importantly, inhibiting sphingosine-1-phosphate (S1P)-dependent egress of lymphocytes from lymphoid tissues with the sphingosine-1-phosphate receptor 1 (S1PR1) agonist FTY720^32^ reduced circulating B cells and Iε-tdTomato^+^ B cells in the lung vasculature significantly but did not reduce the number of Iε-tdTomato^+^ B cells in the lung tissue (**Figure S3**), indicating that Iε-tdTomato^+^ B cells do not traffic to the lungs after reporter activation in lymph nodes but rather undergo IgE CSR *in situ* in lung tissue.

**Figure 2.**
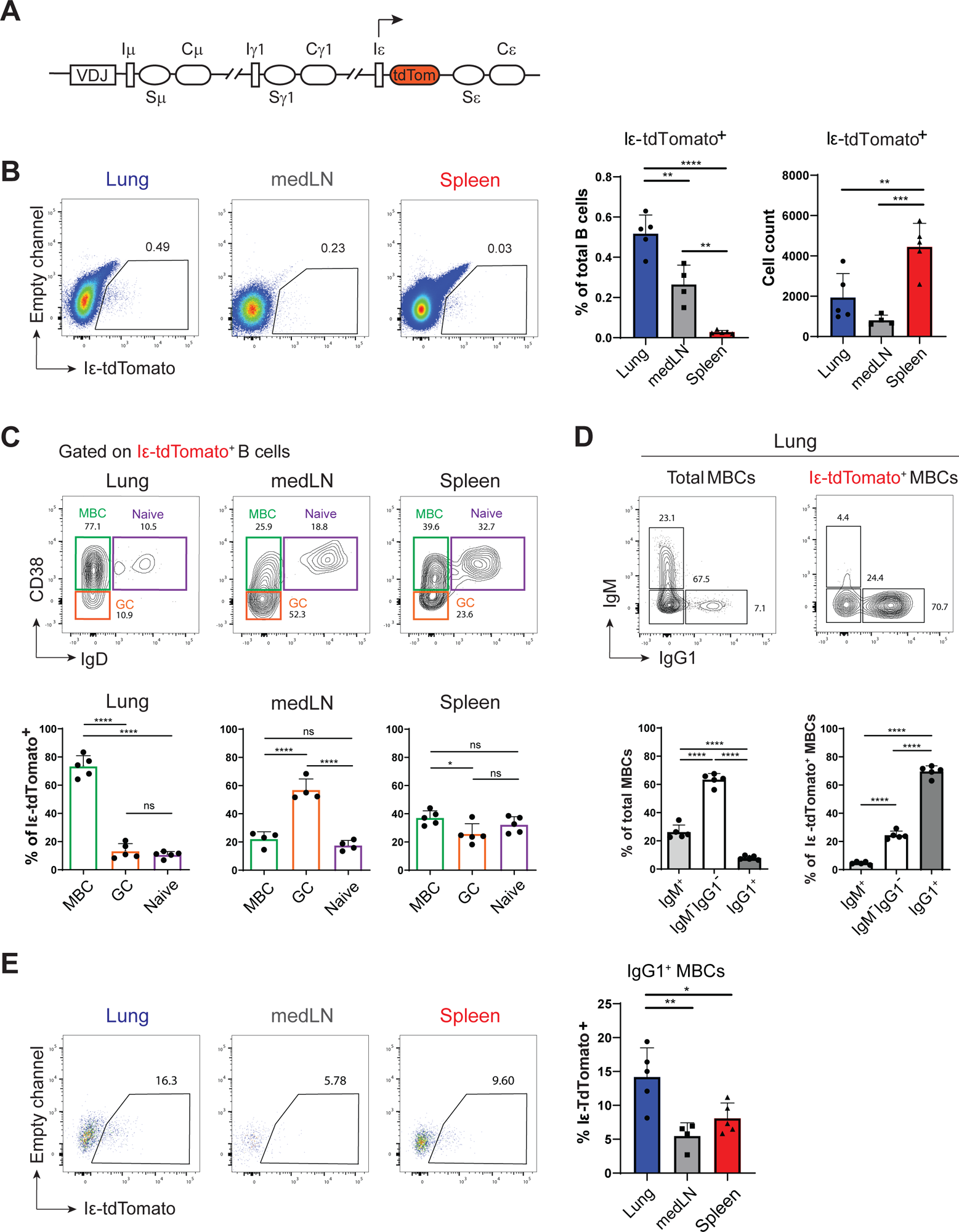
Analysis of IgE class switching using Iε-tdTomato reporter mice. **(A)** Schematic of Iε-tdTomato knock-in mouse reporting germline transcription at Iε locus. **(B)** Representative plots by flow cytometry showing Iε-tdTomato expression by B cells (live B220^+^ i.v. CD45^-^) in the lungs, medLN, and spleen on day 21 (left) and frequencies and cell counts of Iε-tdTomato^+^ B cells (right). **(C)** Representative plots by flow cytometry showing the identification of naïve, germinal center (GC), and memory B cells (MBCs, top) and frequencies of each subset (bottom) among total Iε-tdTomato^+^ B cells. **(D)** Surface expression of IgM and IgG1 on total MBCs and Iε-tdTomato^+^ MBCs in the lungs (top) and frequencies of each population (bottom). **(E)** Expression of Iε-tdTomato among IgG1^+^ MBCs in the lungs, medLN, and spleen. Data are representative of three independent experiments with 5-9 mice in each experiment. One-way ANOVA with Tukey’s multiple comparisons test on mean ± SD; *, p<0.05; **, p<0.01; ***, p<0.001; ****, p<0.0001).

CSR was thought to predominantly occur in germinal centers^33^. However, recent evidence suggests that activated B cells mostly undergo CSR prior to GC formation^34^. Furthermore, MBCs can also undergo CSR during re-activation^35^. To determine which B cell subsets undergo IgE CSR, we analyzed the expression of surface markers on Iε-tdTomato^+^ B cells. In the lungs, the majority of Iε-tdTomato^+^ B cells were MBCs (CD38^+^IgD^-^), with naïve B cells (CD38^+^IgD^+^) and GC B cells (CD38^-^IgD^-^) representing only a minor fraction (∼10%) of IgE-switching B cells (**Figure 2C**). In contrast, GC B cells (CD38^-^IgD^-^) were the major population of IgE-switching B cells in the medLN (**Figure 2C**). In the spleen, IgE-switching B cells were comprised of MBCs, GC B cells, and naïve B cells in similar proportions (**Figure 2C**).

IgE CSR can occur directly from unswitched, IgM^+^ B cells, giving rise to low-affinity IgE, or sequentially via IgG-expressing B cells to generate high-affinity IgE antibodies^36^. To identify the pathway by which IgE-expressing cells arise during allergic airway inflammation, we analyzed the surface expression of IgM and IgG1, the predominant IgG isotype expressed during T_H_2 responses, on Iε-tdTomato^+^ B cells. We observed that most of the Iε-tdTomato^+^ MBCs expressed IgG1, and very few expressed IgM, indicating that sequential switching via IgG1^+^ MBCs is the dominant pathway of IgE CSR in the lung. Strikingly, over 70% of Iε-tdTomato^+^ lung MBCs are IgG1^+^, although IgG1^+^ MBCs represent less than 10% of the total lung MBCs, suggesting an enhanced propensity of IgG1^+^ MBCs to undergo IgE class switching (**Figure 2D**). Furthermore, while IgG1^+^ MBCs could be detected in the lungs and lymphoid organs, IgG1^+^ MBCs in the lungs expressed Iε-tdTomato at a higher frequency than those in the medLN or spleen (**Figure 2E**), indicating that the lungs may be a permissive environment for IgE CSR.

### MBCs and IL-4^+^ T_H_2 cells collaborate to promote local IgE CSR in the lungs

We next investigated the factors that drive MBCs to undergo local IgE CSR in the lungs. IgE CSR depends on cognate interactions with CD4^+^ helper T cells that provide IL-4 and CD40 ligand (CD40L) stimulation to B cells^37,38^. The elevated frequency of IgE-switching B cells in the lungs led us to hypothesize that IL-4-producing CD4^+^ T cells may be abundant in the lungs, therefore creating a permissive environment for IgE CSR. Using immunofluorescence microscopy on lung sections from mice after allergen inhalation, we found that B cells were localized with CD4^+^ T cells in peri-bronchiolar lymphoid aggregates (**Figure 3A**). We compared the frequency of IL-4-producing CD4^+^ T cells in the lungs and lymphoid organs by flow cytometry using IL-4/GFP-enhanced transcript (4Get) mice^39^ and found that a significantly higher frequency of CD4^+^ T cells expressed eGFP in the lungs than the medLN or spleen (**Figure 3B**). T helper type 2 (T_H_2) and T_FH_ cells are major producers of IL-4^40^, and both subsets have been reported to induce IgE responses^41–43^. We detected very few CXCR5^+^PD-1^high^ T_FH_ cells in the lungs despite the presence of GC-phenotype B cells **(Figure S2A)**; however, T_FH_ were present in the medLN and spleen and most expressed eGFP (**Figure S4A**). In contrast, more than 30% of CD4^+^ T cells in the lungs expressed the IL-33 receptor ST2, a marker for T_H_2 cells, while T_H_2 cells were present at low frequencies in the medLN and spleen (**Figure S4B**). Around 50% of eGFP^+^ effector T cells in the lungs were ST2^+^ T_H_2 cells, while only around 20% of eGFP^+^ effector T cells expressed ST2 in the medLN and spleen (**Figure S4C**).

**Figure 3.**
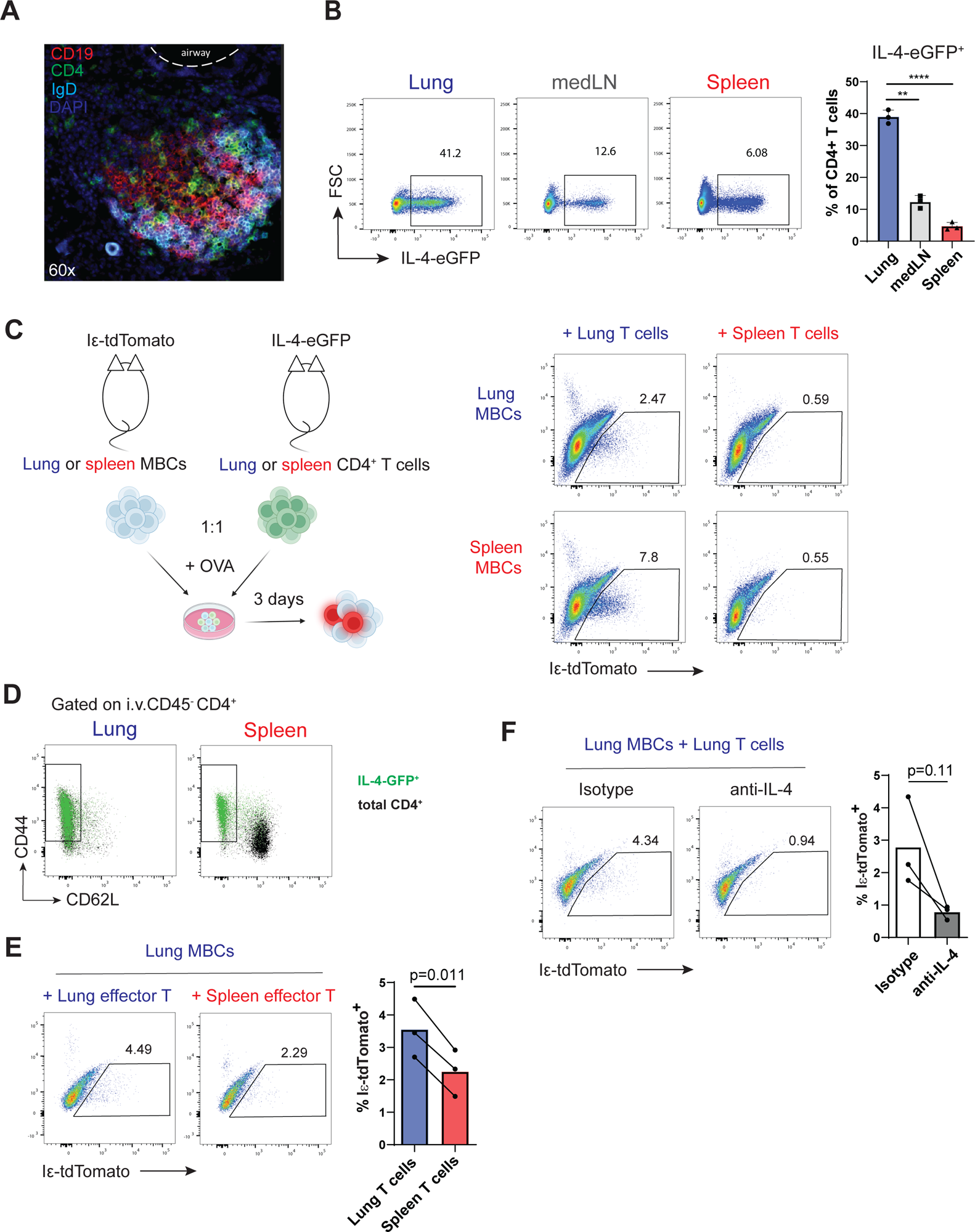
Control of IgE class switching by CD4^+^ T cells. **(A)** Immunofluorescence staining of sensitized lung cryosections harvested on day 21 (60x magnification) showing localization of total B cells (CD19^+^), naïve B cells (IgD^+^), and CD4^+^ T cells (CD4^+^). **(B)** Frequencies of eGFP among CD4^+^ T cells in the lungs, medLN, and spleen of sensitized 4Get mice on day 21. Representative of two independent experiments. **(C)** MBCs were isolated from the lungs or spleen of sensitized Iε-tdTomato reporter mice by FACS and cultured at 1:1 ratio (5×10^4^ of each per well) with CD4^+^ T cells from the lungs or spleen of IL4-eGFP (4Get) mice in the presence of OVA (10 μg/mL) (left) and analyzed on day 3. The frequency of tdTomato^+^ MBCs on day 3 (right). Created with BioRender.com. **(D)** Representative plots of CD44 and CD62L expression on day 21 showing overlay of GFP^+^ (green) CD4^+^ T cells and total CD4^+^ T cells (black). Representative of 2 independent experiments with 3 mice. **(E)** Frequency of tdTomato^+^ MBCs after culture with CD4^+^ CD62L^-^ effector T cells from the lungs or spleen of sensitized mice. Paired *t*-test with lines connecting results from three independent experiments. **(F)** Frequency of tdTomato^+^ MBCs after culture with effector T cells from sensitized mice in the presence of anti-IL4 (10 μg/mL) or isotype control antibody. Paired *t*-test with lines connecting results from three independent experiments.

The scarcity of T_FH_ cells and the abundance of IL-4-eGFP^+^ T_H_2 cells in the lungs led us to hypothesize that cognate interactions between T_H_2 cells and MBCs drive elevated IgE CSR in the lungs. To test this hypothesis, we isolated MBCs from the lungs or spleen of allergen-treated Iε-tdTomato reporter mice and cultured them together with CD4^+^ T cells from the lungs or spleen of allergen-treated 4Get mice in the presence of OVA (**Figure 3C**). After 3 days in culture, we measured the frequency of Iε-tdTomato^+^ B cells and found that IgE switching was elevated in the presence of lung T cells, with minimal Iε-tdTomato^+^ B cells in the presence of spleen T cells (**Figure 3C**). Importantly, expression of Iε-tdTomato was dependent on both antigen and T cells, since minimal Iε-tdTomato expression was observed in conditions without either OVA or T cells (**Figure S5**), indicating the dependency of cognate interaction with T cells in inducing IgE switching. In the lungs and spleen, IL-4-eGFP^+^ CD4^+^ T cells expressed CD44 and lacked CD62L, a phenotype consistent with effector T cells (**Figure 3D**). Additionally, since CD4^+^ T cells in the lungs were almost entirely CD44^+^CD62L^-^ effector T cells, while many CD4^+^ T cells in the spleen were CD44^-^CD62L^+^ naïve T cells (**Figure 3D**), we reasoned that the enrichment of effector T cells may explain the elevated ability of lung T cells to promote IgE CSR. We therefore repeated coculture experiments with effector T cells after depletion of CD62L^+^ T cells. We found that lung effector T cells induced elevated expression of Iε-tdTomato in lung MBCs compared to conditions with spleen effector T cells (**Figure 3E**). Finally, since lung effector T cells expressed higher frequencies of IL-4-eGFP, we neutralized IL-4 in the coculture system with the addition of anti-IL-4 antibodies and observed a reduction of Iε-tdTomato^+^ cells to background levels (**Figure 3F**), indicating that the abundance of IL-4^+^ T_H_2 cells in the lungs drives elevated IgE CSR by lung MBCs.

### Persistence of lymphoid aggregates and residency of lung memory B cells

Peri-bronchiolar lymphoid aggregates containing B cells and CD4^+^ T cells remained in the lungs 8 weeks after allergen inhalation (**Figure 4A**). Since MBCs were principal IgE-switching B cells in the lungs, we asked whether they were maintained in the lungs. We enumerated MBCs immediately (day 21) or 4, 8, or 24 weeks after allergen sensitization, and found that both IgM^+^ and IgG1^+^ MBCs were maintained in the lungs 24 weeks after allergen exposure (**Figure 4B**). To test whether lung MBCs were resident in the lungs or recirculating, we treated mice with FTY720 for 7 days to inhibit sphingosine-1-phosphate (S1P)-dependent lymphocyte recirculation from lymph nodes 8 weeks after allergen inhalation. The numbers of MBCs (i.v. anti-CD45^+^) in the lung vasculature were significantly reduced in mice treated with FTY720 relative to saline controls (**Figure S6A**). However, in the lung tissues (i.v. anti-CD45^-^), MBCs were not reduced while naïve B cells were significantly reduced, indicating that lung MBCs, but not naïve B cells, are stably localized to the lung tissue and not continuously recirculating through the lungs via the blood (**Figure S6A**). However, we considered that FTY720 may also block S1P-dependent egress of MBCs from the lungs into lymphatic vessels, potentially trapping recirculating MBCs in the lung tissue. We therefore injected mice with anti-CD20 antibodies (clone 5D2), which are reported to deplete recirculating B cells^27,44^ (**Figure S6B**). In these experiments, we observed a near total loss of MBCs in the lung vasculature, while MBCs in lung tissues were only partially depleted, suggesting that some lung MBCs are resident in lung tissues. To formally test whether MBCs are resident in lung tissue, we performed parabiosis experiments using congenic CD45.1 and CD45.2 mice (**Figure 4C**). Three weeks after allergen inhalation, mice were surgically paired for 10 days to allow for the equilibration of circulating cells. In the lung vasculature and blood we observed complete chimerism of MBCs with ∼50% of MBCs originating from the host (CD45.2) and partner (CD45.1) mice (**Figures 4D and 4E**). In the spleen, which is known to contain both resident and recirculating MBCs^45^, we observed incomplete chimerism with ∼65% of MBCs remaining host origin (**Figures 4D and 4E**). Strikingly, lung MBCs remained almost entirely host origin, indicating that they are stably resident in lung tissue and minimally recirculate (**Figures 4D and 4E**).

**Figure 4.**
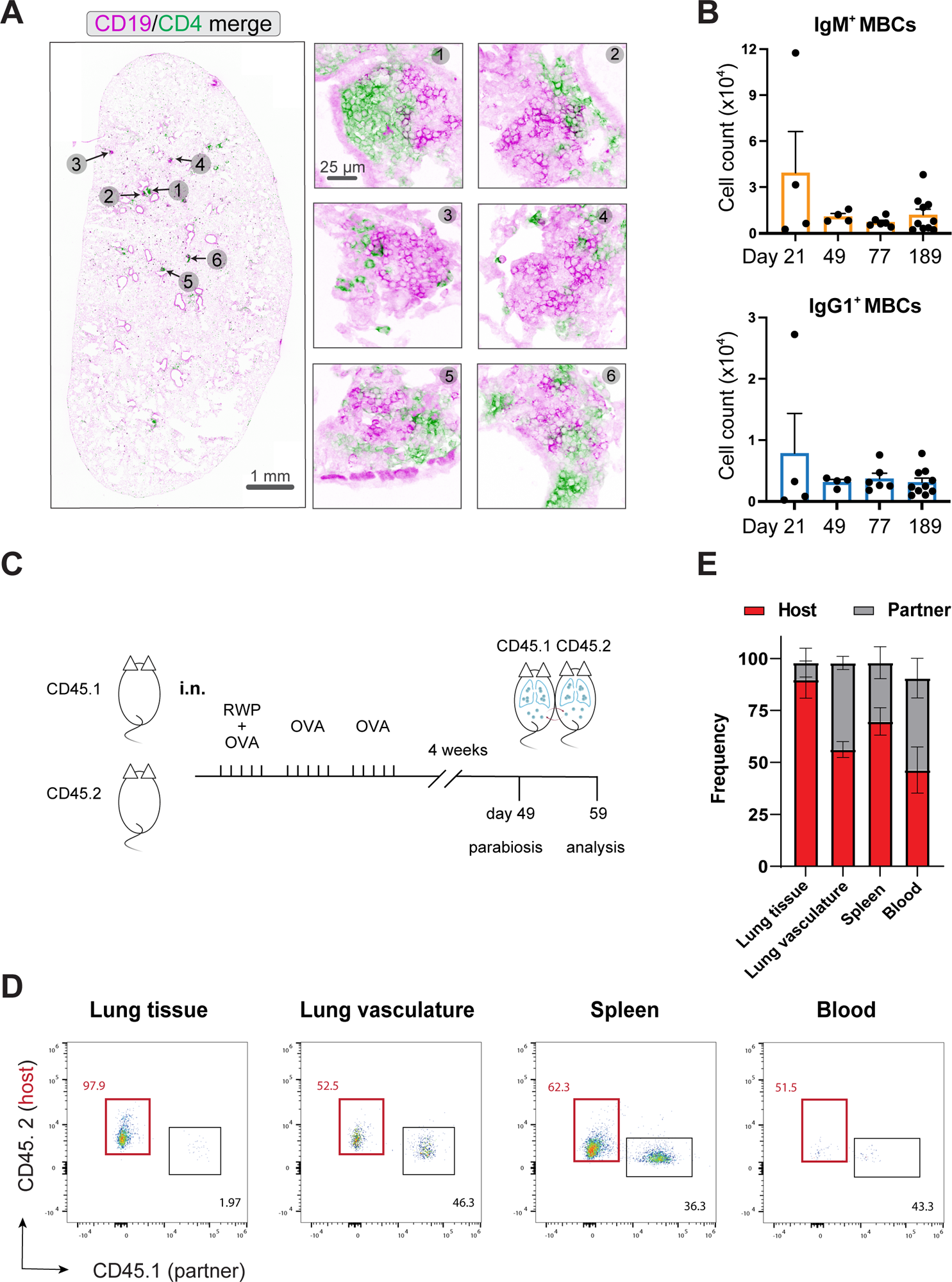
Residency of memory B cells in the lungs. **(A)** Immunofluorescence imaging showing localization of B cells (CD19^+^) and CD4^+^ T cells on day 77. Whole lung section (10x, left) and 63x magnification of indicated areas (right). **(B)** Number of IgM^+^ (top) and IgG1^+^ (bottom) MBCs in the lungs at indicated timepoints. Mean ± SEM, representative of two independent experiments. **(C)** Experimental scheme for parabiosis experiments. CD45 congenic mice were surgically paired 4 weeks after allergen sensitization and tissues were analyzed for MBC chimerism 10 days later. **(D)** Representative flow cytometry plots and **(E)** aggregate frequencies of chimerism of MBCs in indicated tissues based on expression of CD45.1 and CD45.2. Mean ± SD, n=6.

### Persistent lung MBCs are transcriptomically heterogeneous with shared signatures from early lung MBC subsets

MBCs within the lung tissues (i.v. CD45^-^) are tissue-resident. We hypothesized that populations present immediately after sensitization (day 21) are more heterogeneous with dynamic subsets composed of proliferating and differentiating cells, whereas later, when lung inflammation per the measurement of eosinophilia has subsided (day 77), the populations would represent stable lung populations (**Figure S7**). We tested this idea by performing single-cell RNA-seq analysis to profile the heterogeneity of FACS purified MBCs (i.v. CD45^-^B220^+^CD38^+^IgD^-^GL7^-^) from these two time points. Unsupervised clustering analyses and dimensionality reduction analyses by UMAP on the day 21 libraries revealed the presence of eight clusters (Clusters 0 to 7) (**Figure 5A**). Cluster 7 contained only two cells and was therefore excluded from further analyses. Although we imposed filtering criteria to exclude cells of poor quality, marker genes in Cluster 6 included several mitochondrial genes (*Mt-Co1, Mt-Co* and *Mt-Co3*), suggesting that this cluster may be contaminated with dying cells. Marker genes for Cluster 5 included *Ighd* and *Fcer2a*, which resemble phenotypes of follicular B cells (**Figure S8A**). Cluster 4 cells express the marker gene *Irf4*, which suggests this subset may be differentiating into plasma cells. The remaining major clusters 0, 1, 2, and 3 are distinguishable by marker genes including *Plac8, Odc1, Ighm*, and *Ighg1* (**Figure S8A**).

**Figure 5.**
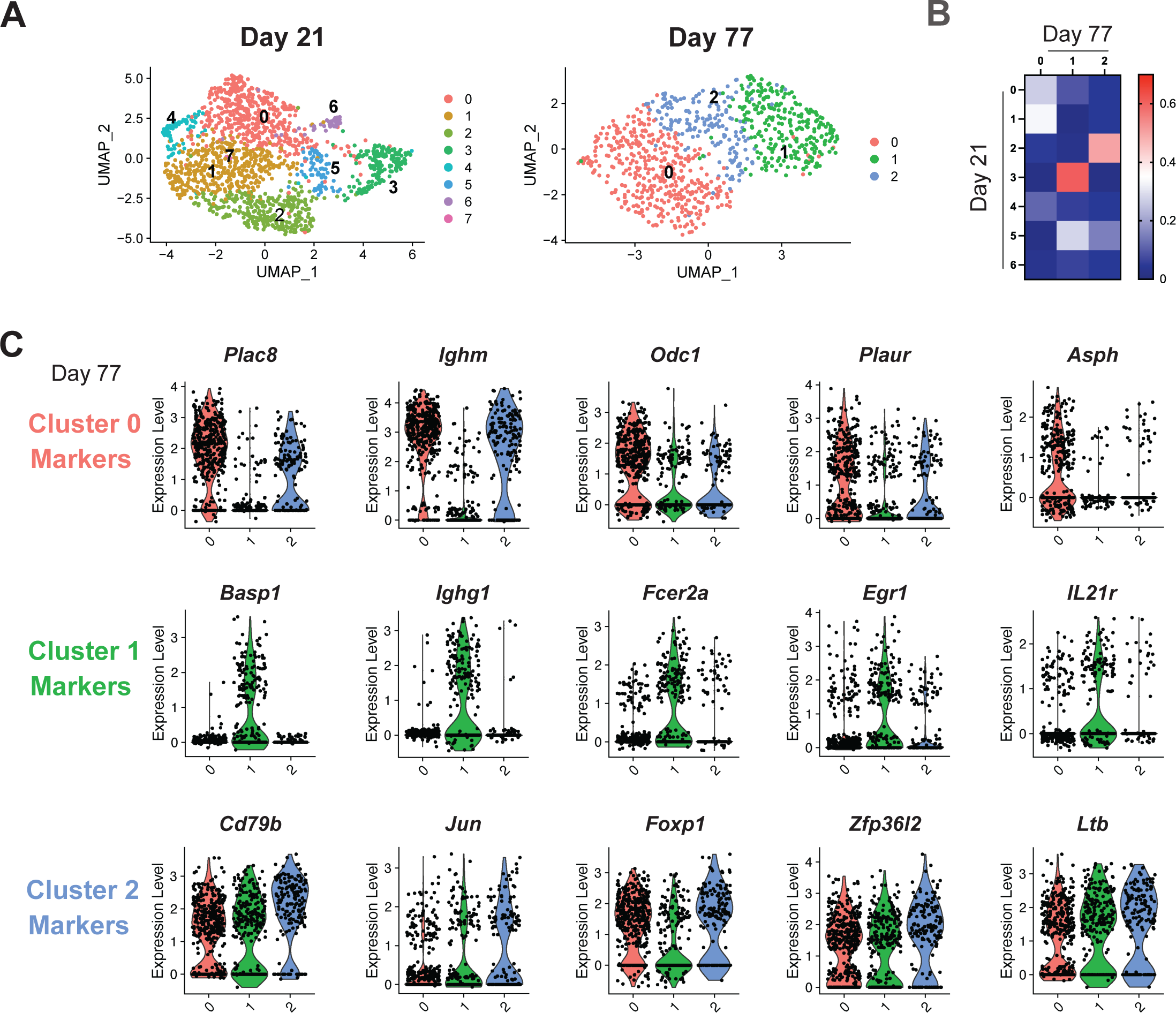
Transcriptomic heterogeneity of lung MBCs. **(A)** UMAP (Uniform Manifold Approximation and Projection) plots showing clusters of lung MBCs on day 21 and 77. Each time point contains two individual libraries. Scale is normalized UMI (Unique Molecular Identifier) counts. **(B)** Szymkiewicz-Simpson similarity matrix showing similarity of marker genes shared between day 21 and day 77 clusters. **(C)** Violin plots showing marker genes from the three clusters in day 77 libraries. Data are scaled expression based on normalized UMI counts.

In contrast, similar analyses on day 77 libraries revealed only three clusters with marker genes that overlap with those for Clusters 0, 1, 2, and 3 on day 21 (**Figures 5A and S8B**). We calculated the Szymkiewicz-Simpson similarity coefficients between clusters present on day 21 and day 77 by analyzing the overlap of marker genes for each cluster (**Figure 5B**). Specifically, day 77 Cluster 0 shared marker genes such as *Plac8* and *Plaur* with day 21 Cluster 0, and marker genes including *Odc1* and *Ighm* with day 21 Cluster 1 (**Figures 5C and S9)**. Day 77 Cluster 1 highly resembled day 21 Cluster 3, sharing marker genes such as *Basp1*, *Ighg1*, *Fcer2a*, *Erg1* and *IL21r* (**Figures 5C and S9**). Day 77 Cluster 2 and day 21 Cluster 2 exhibited similar expression of *Ighm* and shared marker genes for signaling and transcriptional regulation functions including *Cd79a*, *Jun* and *Foxp1* (**Figures 5C and S9**). Altogether, these results indicate that three subsets of lung-infiltrating MBCs with distinct transcriptomic signatures are retained in the lungs after the resolution of inflammation as lung-resident MBCs.

### Phenotypic analyses of lung MBC markers reveal potential mechanisms for their persistency in the lung

The persistence of MBCs in the lung is likely a combined result of recruitment to lung tissues and retention within survival niches in the lungs. Therefore, we probed our single-cell RNA-seq data for the expression of chemokine receptors, adhesion molecules, and potential retention factors. Notably, all three clusters on day 77 expressed *Ccr6, Itga4,* and *Cd69* (**Figure 6A**). Analysis by flow cytometry on day 77 revealed that MBCs in lung tissue expressed higher levels of CCR6 than MBCs in the lung vasculature, medLN, or spleen (**Figure 6B**). Similarly, the integrin CD49d (integrin α4) was expressed on almost all MBCs but at a higher level on MBCs in lung tissue (MFI=20,645 ± 2,577) compared to MBCs in lung vasculature (MFI=10,293 ± 1,543), medLN (MFI=14,755 ± 1,543), or spleen (MFI=6,345 ± 709) **(Figure 6B**). Lung MBCs also expressed the highest levels of CD29 (integrin β1) and CD11a (integrin αL) despite variable detection of the genes encoding these integrins (*Itgal, Itgb1*) among day 77 clusters (**Figures 6A and 6B**). Overall, the surface expression of integrins and CCR6 were markedly different between the lung and lung vasculature, indicating a potential role of these proteins in mediating trafficking and/or retention in lung tissue. Additionally, the surface phenotype of lung MBCs was most similar to MBCs in the medLN (**Figure 6B**), suggesting that MBCs in these two tissues may be related. Finally, expression of CD69 is considered a key marker for tissue-resident T cells^25^, and we observed *Cd69* expression by all three clusters (**Figure 6A**) and surface expression of CD69 on ∼25% of lung MBCs (**Figure 6B**). The differential expression of chemokine receptors and adhesion molecules on MBCs in different tissues suggests that these molecules may be functionally responsible for the lung localization of MBCs.

**Figure 6.**
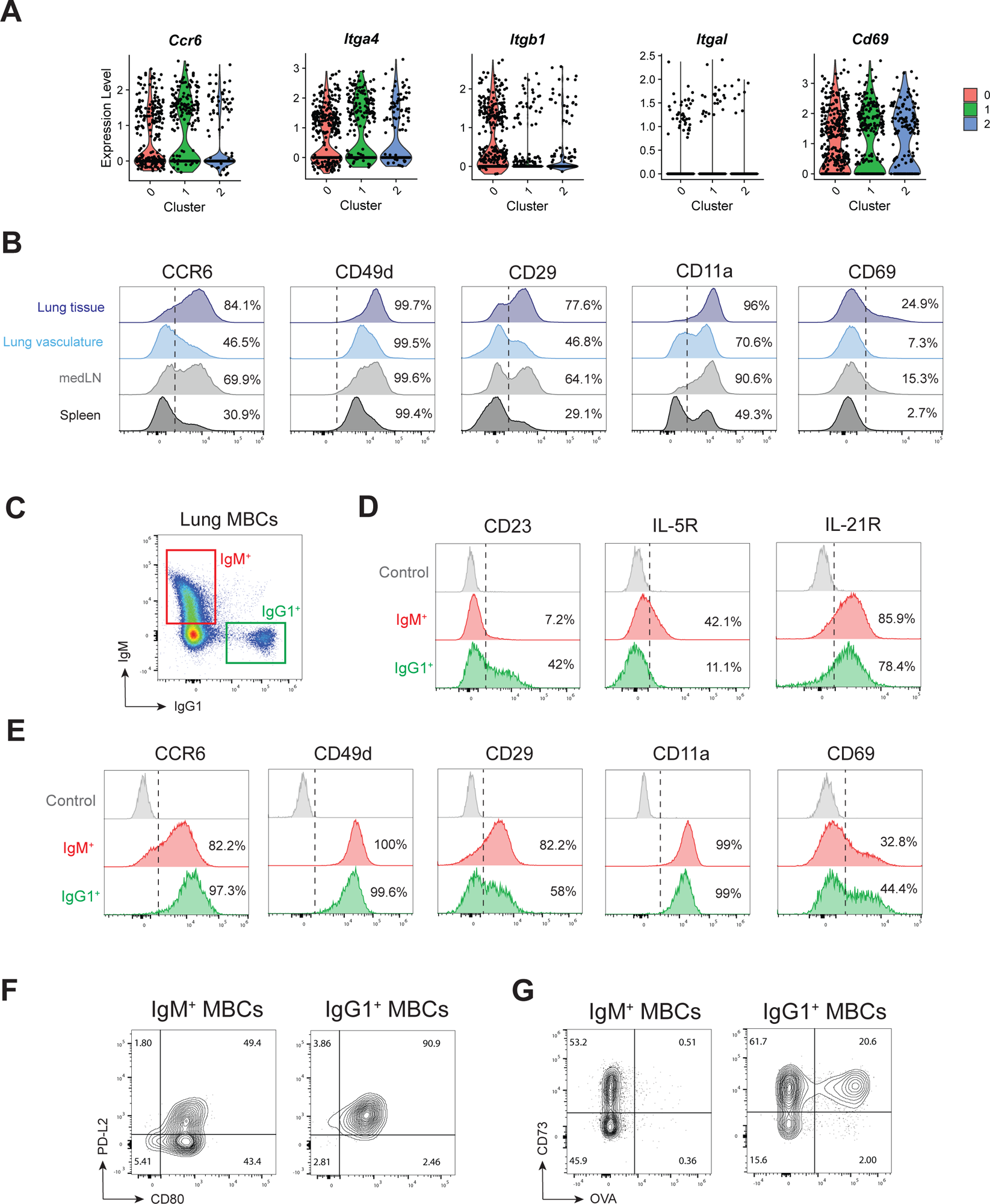
Surface markers of lung memory B cells. **(A)** Violin plots showing scaled expression of indicated genes from day 77 scRNA-seq libraries. **(B)** Histograms showing surface staining of indicated markers on MBCs from each tissue on day 77 by flow cytometry. Each histogram is concatenated from 8 mice. Dashed lines indicate staining from negative control. **(C)** Expression and gating of IgM and IgG1 on MBCs in the lungs (i.v. CD45^-^) on day 77. **(D)** Histograms showing surface expression of markers from day 77 libraries and **(E)** surface markers from A and B on IgM^+^ and IgG1^+^ MBCs. Each histogram is concatenated from 8 mice. Control is only stained with CD19 and live/dead dye zombie NIR. **(E)** Expression of PD-L2 and CD80 and **(G)** CD73 and OVA-binding on IgM^+^ and IgG1^+^ MBCs by flow cytometry on day 77. Representative staining from two independent experiments with 8 mice.

Single-cell RNA-seq analyses on MBCs revealed transcriptomically distinct clusters that are marked by the expression of *Ighg1* and *Ighm* (**Figure 5**), suggesting that the *Ighg1*- and *Ighm*-expressing MBC subsets may possess intrinsic differences in tissue residency and/or function. We therefore profiled molecules that potentially contribute to the lung residency and functions of IgG1^+^ and IgM^+^ MBCs by flow cytometry (**Figure 6C**). Similar to the transcriptional profiles we observed for *Ighm-* and *Ighg1*-expressing MBCs (**Figure 5C**), we observed elevated expression of the IgE Fc receptor CD23 (*FcεR2a*) on IgG1^+^ MBCs and elevated levels of IL-5 receptor (IL-5R) on IgM^+^ MBCs, with both populations expressing similar levels of IL-21 receptor (IL-21R, **Figure 6D**). While IgG1^+^ MBCs expressed higher levels of CCR6 and IgM^+^ MBCs expressed more CD29, both IgM^+^ and IgG1^+^ MBCs expressed high levels of CD11a and CD49d (**Figure 6E**). A portion of IgM^+^ and IgG1^+^ MBCs also expressed CD69 (∼33% and 44%, respectively, **Figure 6E**). Previous studies have demonstrated that subsets of MBCs with different functions can be identified based on expression of CD80, CD73, and PD-L2^46,47^. Approximately 50% of IgM^+^ MBCs expressed CD80 and PD-L2, while IgG1^+^ MBCs in the lungs were uniformly CD80^+^PD-L2^+^ (**Figures 6F and 6G**), a phenotype associated with an intrinsic propensity toward plasma cell fate^46^ (**Figure 6F**). Similarly, while only around 50% of IgM^+^ MBCs were CD73^+^, the majority of IgG1^+^ MBCs also expressed CD73, a marker of GC experience^48,49^ (**Figure 6G**). Staining for OVA-binding B cells by flow cytometry revealed that IgG1^+^, but not IgM^+^, MBCs exhibited OVA-specificity (**Figure 6G**), suggesting that at least some IgG1^+^ MBCs were generated from GC reactions where affinity maturation towards OVA occurred.

### Function of lung memory B cells during recall response

We found that lung MBCs are the predominant IgE switching cells that persist in the lung with stable transcriptomic signatures. Therefore, we hypothesized that lung-resident MBCs may be a reservoir for IgE responses upon allergen re-exposure in the respiratory tract. To determine if lung-resident MBCs mediate local IgE responses to allergen re-encounter, we modeled allergen exposure by nebulization mice with OVA (**Figure 7A**). Rechallenge with OVA induced a robust increase in OVA-specific IgE in BALF (**Figure 7B**), formation of OVA-specific IgE-secreting cells in the lungs (**Figure 7C**), and expansion of OVA-specific MBCs in the lungs (**Figures 7D and 7E**). These recall responses depend on antigen-specific MBCs because unsensitized mice nebulized with OVA failed to produce detectable local OVA-specific IgE or OVA-specific IgE-secreting cells (**Figures 7B and 7C**). Therefore, local IgE responses to OVA rechallenge depend on the activation of OVA-specific MBCs rather than *de novo* formation of IgE-producing B cells. Additionally, IgE responses are primarily localized to the airway since OVA-specific IgE-secreting cells were abundant in the lungs but not observed in the bone marrow (**Figure 7C**). In contrast, OVA-specific IgG1-secreting cells were abundant in the lung, spleen, and bone marrow (**Figure S10**).

**Figure 7.**
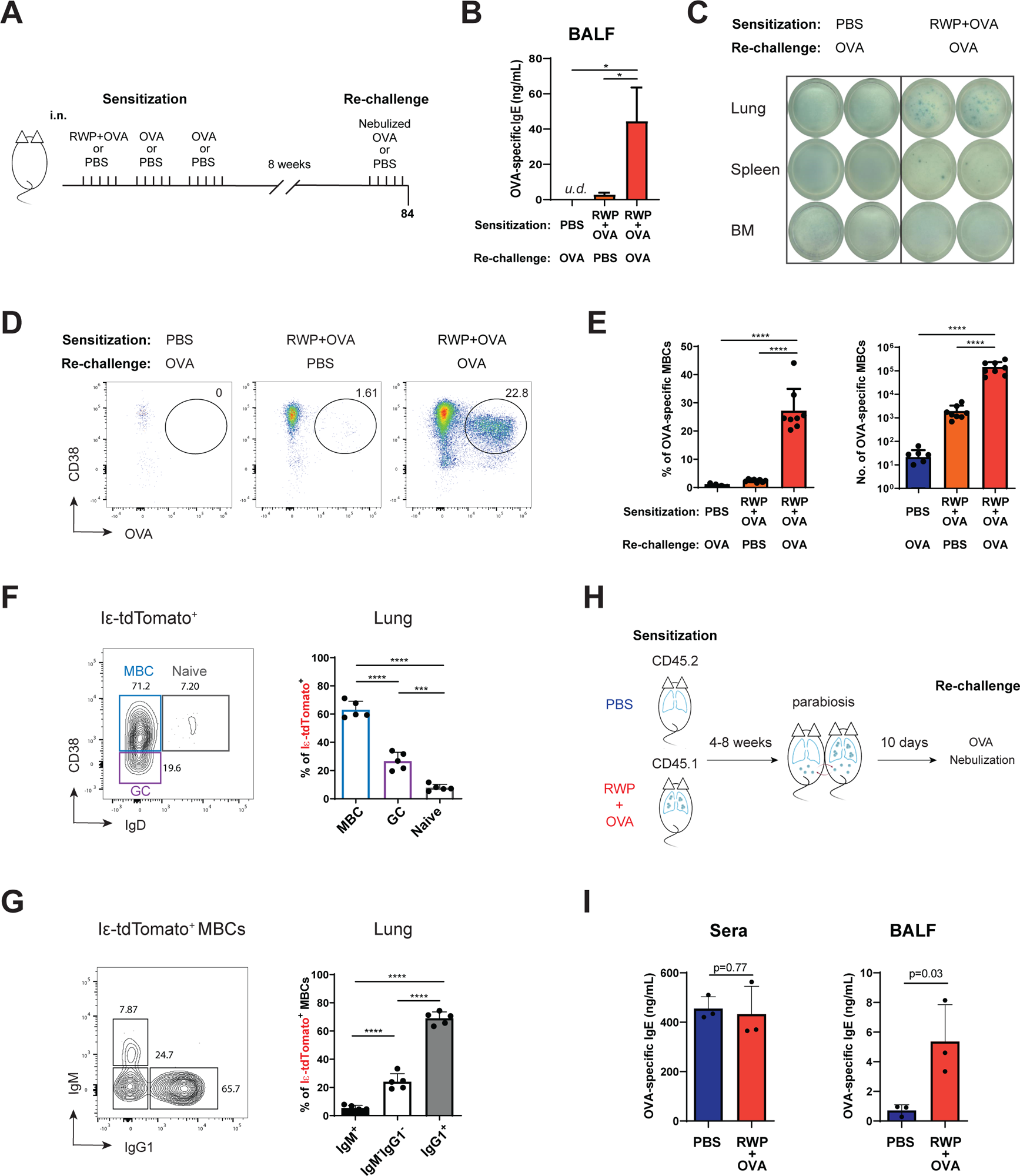
Function of lung-resident MBCs during recall response. **(A)** Experimental schematic of rechallenge. Eight weeks after sensitization with RWP+OVA (or PBS control) mice were rechallenged with five consecutive days of nebulized OVA (in 1% PBS) or PBS (20 min/day) and analyzed on day 84. **(B)** Detection of OVA-specific IgE in bronchoalveolar lavage fluid (BALF) by ELISA. One-way ANOVA with Tukey’s multiple comparisons test (n=8). **(C)** Detection of OVA-specific IgE-secreting cells in single cell suspension from lungs, spleen or bone marrow (BM) by ELISpot with 1×10^6^ cells plated per well. **(D)** Representative flow cytometry plots showing the identification of OVA-binding MBCs (i.v. CD45^-^ CD19^+^ IgD^-^) in the lungs in each indicated condition. **(E)** Frequencies and numbers of OVA-specific MBCs in the lungs in each indicated condition. One-way ANOVA with Tukey’s multiple comparison’s test. **(F)** Representative (left) and summary plot (right) showing frequency of MBCs, naïve, and germinal center (GC) B cells among Iε-tdTomato^+^ B cells in the lungs. Representative of three independent experiments, one-way ANOVA with Tukey’s multiple comparison’s test. **(G)** Expression of IgM and IgG1 among Iε-tdTomato^+^ MBCs in the lungs. Representative plot (left) and aggregate frequencies by flow cytometry analyses (right). Representative of three independent experiments, one-way ANOVA with Tukey’s multiple comparison’s test. **(H)** Schematic of parabiosis experiments. RWP+OVA-sensitized mice were surgically paired to PBS control mice 4 or 8 weeks after sensitization. Ten days after surgery, pairs were rechallenged with five days of nebulized OVA (1% in PBS, 20 min/day). **(I)** Detection of OVA-specific IgE in sera and BALF of PBS or RWP+OVA-sensitized parabionts by ELISA after rechallenge with OVA. Representative data of two independent experiments. Bars are mean ± SD; *, p<0.05; ***, p<0.001; ****, p<0.0001.

Similar to primary sensitization with allergen, IgE-switching (Iε-tdTomato^+^) B cells in the lungs were mostly (63±5%) MBCs (**Figure 7F**), and most (69±4%) of these MBCs expressed IgG1 (**Figure 7G**), indicating that IgE responses in the airway depend on re-activation of MBCs that undergo sequential CSR to IgE in the lungs. To determine whether lung-resident MBCs drive the production of IgE in the airway, we surgically paired allergen-sensitized (RWP+OVA) or unsensitized (PBS) mice and rechallenged parabionts with nebulized OVA (**Figure 7H**). OVA-specific IgE levels were equal in sera of both parabionts; however, allergen-sensitized (RWP+OVA) mice with lung-resident MBCs had significantly more OVA-specific IgE in the airway (**Figure 7I**), indicating that circulating IgE contributes minimally to airway IgE. Altogether, these data demonstrate that lung-resident MBCs drive local IgE responses in the airway.

## DISCUSSION

The lack of IgE^+^ memory cells and the absence of systemic IgE responses in some allergy patients are at odds with the chronic nature of airway hypersensitivity in allergic asthma. In this study, we sought to identify the tissue sites and cell types that support IgE class switching. Using a mouse model based on local allergen exposure in the airway and reporter mice to identify IgE-switching B cells, we found that the sensitized lungs were a major site of IgE production. Allergen inhalation induced B cell infiltration into the lungs, including IgM^+^ and IgG1^+^ MBCs. Most IgE-switching B cells in the lungs were IgG1^+^ MBCs, indicating that sequential CSR via IgG1^+^ MBCs is the predominant pathway of IgE production in the respiratory tract. MBCs were maintained in the lungs for at least 6 months and parabiosis experiments revealed that MBCs were resident in lung tissue, suggesting they may be a reservoir for allergen-specific IgE in the respiratory tract. To test this possibility, we rechallenged mice with antigen and observed expansion of antigen-specific MBCs, increases in antigen-specific IgE in the airway, and formation of antigen-specific IgE-secreting cells in the lungs. Furthermore, rechallenge of allergen-sensitized mice surgically paired to non-sensitized mice induced robust production of IgE only in the airway of parabionts with lung-resident MBCs, despite equal levels of circulating IgE in sera. These results indicate that lung-resident MBCs mediate IgE responses in the airway, and that such local IgE responses are physiologically distinct from systemic IgE responses measured by serum IgE.

Adoptively transferred IgG1^+^ MBCs were previously shown to give rise to antigen-specific IgE-producing cells upon antigen challenge, illustrating the ability of IgG1^+^ MBCs to maintain the memory of IgE responses^50^. We identified IgG1^+^ MBCs as the major IgE-switching cells in allergic lung tissues, providing mechanistic evidence for how IgG1^+^ MBCs generate IgE-producing cells. In the same report, the IgG1^+^ MBCs that gave rise to IgE with high affinity to antigen expressed pro-plasma cell markers (PD-L2, CD80 and CD73). We found that over 80% of IgG1^+^ MBCs in the allergic airway express these markers. Additionally, two recent human studies describing an IgG^+^ MBC subset expressing CD23 and IL-4R as well as *IGHE* transcripts were elevated in patients with allergic diseases including allergic asthma^51,52^. Both studies suggested the pathogenic involvement of these IgG^+^ MBCs in allergic diseases as precursors for allergen-specific IgE-producing cells. Our single-cell RNA-seq and phenotypic analyses revealed that IgG1^+^ MBCs in allergic lungs also express high levels of CD23. Altogether, our findings suggest that persistent IgG1^+^ MBCs in the lungs can readily undergo CSR to IgE and rapidly differentiate into allergen-specific IgE-producing cells.

We also found that IgE-switching B cells, mostly MBCs, were more frequent in the lungs than in secondary lymphoid organs, suggesting that allergen-sensitized lungs are a permissive environment for IgE responses. B cells were localized in the lungs together with CD4^+^ T cells in peribronchiolar lymphoid aggregates. CD4^+^ T cells are key drivers of IgE CSR by providing IL-4 and CD40L stimulation to B cells^53,54^. However, the precise T cell subsets that support IgE production are controversial. Data from multiple studies suggested that T follicular helper (T_FH_) cells were critical in generating allergen-specific IgE^42,55,56^. Those studies are often interpreted as T_FH_ directly inducing CSR to IgE. However, the deletion of T_FH_ cells disrupted the proper formation of GCs and thus the output of MBCs and all antigen-specific antibodies including IgG1 were decreased. With the decreased output of MBCs and all antigen-specific antibodies, it is difficult to conclude that T_FH_ cells are required for IgE responses by directly inducing IgE CSR in germinal centers. Additionally, T_FH_ cells that express IL-21 or neuritin have been shown to inhibit IgE responses^57,58^, highlighting the existence of regulatory mechanisms in the germinal center to restrict IgE production. Furthermore, the cell types and mechanisms that support mucosal IgE responses have not been elucidated. In this study, we found that IgE CSR was elevated in the lungs despite the lack of appreciable numbers of T_FH_. Using IL-4 reporter mice, we found that the lungs were enriched in IL-4-producing T cells, which were mostly T_H_2 cells (CD44^+^CD62L^-^ST2^+^). In co-culture experiments with MBCs and effector CD4^+^ T cells, we demonstrated that lung CD4^+^ T cells effectively induced IgE CSR in an antigen and IL-4-dependent manner. We conclude that these lung T_H_2 cells are the source of IL-4 needed for inducing IgE CSR.

We found that the IgE-switching B cell subsets were also drastically different between the sensitized lungs and the mediastinal lymph nodes. In the lungs, IgE CSR was predominately mediated by MBCs, while in the medLNs, GC B cells were the major IgE-switching cells. Although we did not directly test the origin of lung MBCs, IgG1^+^ MBCs in the lungs expressed markers of GC experience (PD-L2, CD80, CD73) and exhibited specificity for OVA, suggesting they were formed in GC responses in the medLN prior to trafficking to the lungs where abundant IL-4^+^ T_H_2 cells induced IgE CSR. Together with differences in the composition of T cell subsets between the lungs and the medLNs, these findings indicate differential regulation of IgE CSR between extrafollicular compartments and the germinal center. Altogether, we propose that IgE CSR in the sensitized lung is driven by the abundance of IL-4-producing CD4^+^ T cells and potentially the lack of T_FH_ subsets with regulatory functions capable of limiting IgE CSR. Additionally, our results demonstrating that IgE CSR was mediated by MBCs outside of the GC is consistent with reports that suggested IgE responses arise via extrafollicular CSR and with the recent realization that CSR does not take place in the germinal centers as frequently as previously thought^34,59^.

Recent studies demonstrate that respiratory infections induce the formation of lung-resident MBCs, which mediate protective local antibody responses to subsequent pathogen encounters^26,27^. In this study, we demonstrated that allergen exposure also elicits the formation of lung-resident MBCs; however, rather than promoting protective antibody responses to pathogens, MBCs in allergic lungs mediate local IgE responses that may drive diseases such as allergic asthma. The mechanisms that mediate MBC trafficking to the lungs and their retention and survival within lung tissue are unknown. We found that allergen inhalation induced the formation of peri-bronchiolar lymphoid aggregates containing B cells and CD4^+^ T cells. These structures may serve as survival niches for MBCs in the lungs, which may be regulated by lung stromal cells that organize these structures by producing chemokines to recruit lymphocytes and provide retention and/or survival signals to MBCs. Lymphocyte entry into tissues is controlled by integrins and chemokine receptors. In our study, we found that lung MBCs express high levels of the integrins CD49d (integrin α4) and CD29 (integrin β1). Heterodimers of CD49d and CD29 (integrin α4β1), also known as VLA-4, bind to vascular cell adhesion molecule-1 (VCAM-1) expressed by vascular endothelial cells during inflammation^60^, raising the possibility that VLA-4 expression by circulating MBCs controls entry into inflamed lungs across VCAM-1^+^ blood endothelial cells. Similar to tissue-resident T cells and recent descriptions of lung-resident MBCs, we observed expression of CD69 on some lung MBCs, albeit at lower frequencies than described after respiratory infection^26,27,61^. Surface expression of CD69 antagonizes sphingosine 1-phosphate receptor-1 (S1PR1), which is proposed to maintain tissue residency by preventing egress from non-lymphoid tissues in response to sphingosine 1-phosphate (S1P)^62,63^. However, CD69 expression is not required for tissue-residency of T cells^64^, and since we observed CD69 expression was only expressed by some lung MBCs, other mechanisms may control the retention of MBCs in the lungs. Physical adhesion to lung stromal cells via CD11a/ICAM-1 interaction may also promote retention and/or survival of MBCs within lymphoid structures in the lungs. We found that lung MBCs express elevated levels of CCR6, which may control the localization of MBCs to the lungs. Lung-resident MBCs induced by respiratory infection also express CCR6^61,65^, suggesting a common pathway by which MBC traffic to the respiratory tract during different types of inflammation. Lung-resident MBCs induced by viral infections express the IFN-γ-inducible chemokine receptor CXCR3^26,28,65^, which we did not detect in our scRNA-seq libraries likely due to the lack of IFN-γ expression after allergen exposure. Therefore, it is likely that different types of inflammation recruit MBCs to the lung mucosa by distinct mechanisms. Understanding these mechanisms will be critical to the development of strategies to prevent the accumulation of MBCs in asthmatic lungs while simultaneously maintaining mucosal immunity to respiratory pathogens.

In summary, this study revealed for the first time the accumulation of lung-resident MBCs in T_H_2 inflammation. These results also provide a mechanistic explanation for how allergen-specific IgE responses arise in the respiratory mucosa even in the absence of systemic IgE. IgG1^+^ memory cells persist in the lungs and form IgE-secreting cells upon re-exposure to allergen. Thus, targeting the formation or accumulation of pathogenic lung-resident MBCs will be important in treating allergic lung inflammation.

## EXPERIMENTAL PROCEDURES

### Experimental Models and Subject Details

#### Mice

All mice were housed in the Comparative Medicine Facility at Loyola University Chicago in specific pathogen-free (SPF) conditions under standard 12:12 hours light/dark cycles. BALB/c, BALB/cByJ, BALB/cByJ CD45.1 congenic mice (CByJ.SJL(B6)-Ptprca/J) and IL-4/GFP-enhanced transcript (4Get, C.129-Il4tm1Lky/J) mice were purchased from Jackson Labs or bred in-house. Iε-tdTomato mice were bred and maintained in-house. All procedures were approved by Loyola University Chicago Institutional Animal Care and Use Committee. Female mice aged 6-10 weeks were used in the *in vivo* experiments. Iε-tdTomato and 4Get mice both males and females aged 6-12 weeks old were sensitized for the isolation of MBCs and T cells in the co-culture experiments.

### Method Details

#### In vivo treatment and immunization

For allergen sensitization 6-10-week-old mice were immunized intranasally with 40 μL of allergen suspension containing 300 μg ragweed pollen (RWP, Greer) and 20 μg ovalbumin (OVA, Sigma Aldrich) in phosphate buffered saline (PBS) for 5 consecutive days in the first week, followed by 2 weeks of 5 consecutive days OVA alone (20 μg) in PBS, under isoflurane anesthesia. Control mice received vehicle (PBS) alone. Mice were analyzed at indicated timepoints after sensitization or rechallenged 8 weeks later. For rechallenge, mice were nebulized with 1% OVA in PBS or PBS alone for 20 minutes per day for five consecutive days in a custom-made chamber and analyzed on day 84. To inhibit lymphocyte egress from lymph nodes the S1PR1 agonist FTY720 was dissolved in 0.9% NaCl and administered daily i.p. at 0.5 mg/kg for 3 or 7 days as indicated. Depletion of circulating B cells was performed using a single i.v. injection of 50 μg anti-mouse CD20 (clone 5D2, a generous gift from Genentech) or IgG2a isotype control (Bio X Cell) in 100 μL PBS by tail vein. Depletion was verified by analysis of numbers of B cells in the blood, lymph nodes, and spleen by flow cytometry. Intravascular labeling of blood immune cells was performed by i.v. injection of 2.4 μg anti-CD45 (conjugated to Pacific Blue or Brilliant Violet 510, BioLegend) in 100 μL PBS by tail vein 3-5 minutes prior to sacrifice.

### Sample preparation and flow cytometry

Mice were euthanized by CO_2_ asphyxiation and cervical dislocation and lungs were perfused with 30 mL of PBS solution containing heparin (10 U/mL, Millipore) via cardiac puncture in the right ventricle while lungs were inflated with air by syringe via catheterization of the trachea. Airway cells were collected by bronchoalveolar lavage by inflating lungs with 1 mL of cold PBS via tracheal catheter and collecting solution into microcentrifuge tubes. Lungs were harvested, then minced by scissors and digested in 2 mL of solution containing collagenase D (750 U/mL, Gibco) and DNase I (200 U/mL, Sigma-Aldrich) in RPMI 1640 (Corning) for 45 minutes shaking at 37 ⁰C. Digested lungs were passed through 70-μm cell strainers and diluted with 10 mL of 30% Percoll (Cytiva) in RPMI, then centrifuged at 930 *g* for 10 minutes at 4⁰ C with no brake. Pellets were resuspended in 5 mL of ACK lysis buffer (150 mM NH4Cl, 10 mM KHCO3, and 0.1 mM EDTA in autoclaved water) for 5 minutes at room temperature (RT) to lyse red blood cells (RBCs). Lysed suspensions were diluted in 10 mL of FACS buffer (PBS with 2% heat inactivated FBS) and centrifuged for at 300 *g* for 5 minutes at 4⁰ C, followed by resuspension in FACS buffer and filtering dead cells through 70-µm cell strainer. Spleens and lymph nodes were collected in PBS and single cell suspensions were prepared by mushing through 70-µm cell strainers, followed by washing and resuspension in FACS buffer. Spleens RBCs were also lysed using ACK lysis buffer (5 mL, 5 minutes, RT). Blood cells were collected by cardiac puncture, followed by lysis of RBCs in 10 mL of ACK lysis buffer, followed by washing with PBS and resuspension in FACS buffer. Cells were counted in 0.4% Trypan Blue using Countess II (Thermo Fisher Scientific).

Isolated cells were washed in FACS buffer and F_C_ receptors were blocked with anti-mouse CD16/32 (Bio X Cell, clone 2.4G2) in FACS buffer for 10 minutes on ice. Dead cells were identified by staining with Zombie Near Infrared Fixable Viability Dye (Thermo Fisher Scientific, 1:2000) for 10 minutes at room temperature (RT) in PBS prior to surface staining, or by addition of DAPI (1:2000 in FACS buffer) after cell staining. Surface staining was performed on ice in 96-well U-bottom plates for 30 minutes using antibodies described in key resources table, and washes were 300 *g* for 5 minutes at 4⁰ C. For detection of OVA-specific B cells, we prepared biotinylated OVA using EZ-link Sulfo-NHS-LC-Biotinylation Kit (Thermo Fisher Scientific), followed by dialysis in PBS for 24 hours to remove free biotin, and storage at 4⁰ C. Biotinylated OVA was added during surface staining, then after 2 washes cells were incubated with streptavidin (SA) conjugated to Brilliant Violet 650 (BV650, 1:200, BioLegend) in PBS on ice for 15 minutes, followed by 3 washes. Similarly, staining for CXCR5 required biotinylated anti-CXCR5 followed by secondary staining with SA-BV650. Cells were acquired on BD Fortessa LSR II or Cytek Aurora 5L using FACSDiva (BD) or SpectroFLo (Cytek) in the FACS Core Facility at Loyola University Chicago. Data were analyzed using FlowJo v10 (Treestar).

### Cell sorting and isolation

For sorting, cells were stained as described above and sorted on a BD FACSAria III into complete media. Memory B cells were sorted as live CD19^+^ or B220^+^ and CD38^+^ IgD^-^ GL7^-^ in addition to gating on intravascular CD45^-^ cells. Total CD4^+^ T cells were sorted as live intravascular CD45^-^ CD3^+^CD4^+^ lymphocytes. Effector CD4^+^ T cells were isolated by magnetic enrichment using EasySep mouse CD4^+^ T cell Isolation Kit (STEMCELL Technologies) with addition of 10 µL of biotinylated anti-CD62L (BioLegend) to remove naïve T cells. Purity was assessed by staining for CD3, CD4, CD44, and CD62L and analysis on Fortessa LSR II.

### Cell culture

Memory B cells and CD4^+^ T cells were resuspended in complete media (Corning RPMI 1640 with L-glutamine plus 10% heat inactivated FBS, 100 U/mL penicillin and 100 µg/mL streptomycin, Corning) and 5×10^4^ of each were combined in 96-well U-bottom plates with anti-CD40 (100 ng/mL, clone HM40-3, BD Biosciences) in the presence or absence of OVA (10 μg/mL, Sigma Aldrich). Cells were cultured for 3 days at 37⁰ C at 5% CO_2_. IL-4 neutralization was performed using anti-IL-4 (10 µg/mL, clone 11B11, BioLegend) or rat IgG1 isotype control (10 µg/mL, clone RTK2071, BioLegend).

### ELISA

Bronchoalveolar lavage fluid (BALF) was collected from lungs at sacrifice and cells removed by pelleting at 700 *g* for 5 minutes and stored at -80⁰ C. Serum was harvested by collecting blood via cardiac puncture, allowing clotting for 2 hours at RT, and centrifuging for 10 minutes (2000 *g*, 4⁰ C) and storage at -80⁰ C. 96-well flat bottom MaxiSorp ELISA plates (Thermo Fisher Scientific) were coated overnight at 4⁰ C in 0.5 M sodium carbonate buffer. For IgE ELISAs, plates were coated with 2 µg/mL of anti-IgE (clone RME-1, BioLegend) and for IgG1 ELISAs, plates were coated with 10 mg/mL of chicken egg ovalbumin (OVA, Sigma-Aldrich). Plates were washed 5 times with 200 µL PBS + 0.05% Tween 20 (PBST) then blocked in 3% bovine serum albumin (BSA, Fisher) for 3 hours at RT. Samples or standards were diluted in 10% FBS in PBS and incubated overnight at 4⁰ C. Sera were diluted 1:10 and 1:50 for IgE, and 1:20,000 and 1:200,000 for IgG1. BALF was added as undiluted or diluted 1:10 for IgE, and 1:500 and 1:5000 for IgG1. Standard curves were constructed by 2-fold dilutions of OVA-specific IgE monoclonal antibody (clone E-C1, Chondrex) diluted in 10% FBS in PBS from 200 ng/mL to 0.195 ng/mL, or with OVA-specific IgG1 antibody (clone L71, Chondrex) from 20 ng/mL to 19.5 pg/mL. Wells were washed 5 times with 200 µL of PBST and 100 µL of diluted detection antibodies were added for 1 hour at RT. For IgE, biotinylated OVA was prepared in-house (see Flow Cytometry section) and diluted 1:1000 in 10% FBS in PBS. For IgG1, biotinylated anti-IgG1 (clone A85-1, BD Biosciences) was added at 1 µg/mL in 10% FBS in PBS. Wells were washed 5 times with 200 µL PBST and streptavidin-conjugated to horseradish peroxidase was added (1:2000 in PBS, BioLegend) for 30 minutes at RT. Wells were washed 5 times with 200 µL PBS and developed by adding 100 µL of 1-Step Turbo TMB ELISA (Thermo Fisher Scientific) for 30 minutes at RT followed by stopping the reaction with 50 µL of 2M H_2_SO_4_ and reading optical density at 450 nm using BioTek ELx800 plate reader.

### ELISpot

ELISpot assays for OVA-specific IgE and IgG1 antibody-secreting cells (ASCs) were performed using Multiscreen white 96-well filter plates with Immobilon-P PVDF membranes (Millipore). Plates were prepared by pre-wetting with 30 µL of 35% ethanol in water for 1 minute, followed by 3 washes with 150 µL of sterile PBS. For OVA-specific IgE assays, plates were coated per well with 100 µL of anti-IgE capture antibody (2 µg/mL in PBS, clone RME-1, BioLegend) overnight at 4⁰ C. For OVA-specific IgG1 assays, plates were coated per well with 100 µL of OVA in PBS (5 µg/mL, Sigma-Aldrich) overnight at 4⁰ C. Plates were washed 4 times with 150 µL of PBS + 0.05% Tween 20 (PBST), then blocked with 100 µL of 20% FBS in RPMI (Hyclone or Corning) per well for 3 hours at 37⁰ C. Cell suspensions from lungs, spleen, or bone marrow were prepared in cell culture RPMI and plated in 100 µL of medium containing 1-4×10^6^ cells or 4-fold serial dilutions in duplicate. As a control for background, the bottom row of plates received medium alone. For OVA-specific IgE, plates were incubated for 24 hours at 37⁰ C in 5% CO2, and for OVA-specific IgG1, plates were incubated for 18 hours. Cells were then removed by dumping medium and washing 5 times with 150 µL of PBST. For detection of OVA-specific IgE ASCs, 100 µL of biotinylated OVA (see Flow Cytometry section) was diluted in PBS with 2% FBS (1:2000 or ∼5 µg/mL) and incubated for 2 hours at RT. For detection of OVA-specific IgG1, wells were incubated with 100 µL of biotinylated anti-IgG1 (1 µg/mL in PBS with 2% FBS, clone A85-1, BD Biosciences) for 2 hours at RT. Plates were washed 5 times in 150 µL of PBST and incubated with 100 µL of streptavidin conjugated to horseradish peroxidase (HRP, BioLegend) diluted 1:2000 in PBS for 1 hour at RT, followed by 5 washes with 150 µL of PBS. Spots were developed by adding 100 µL of TMB (3,3’,5,5’-Tetramethylbenzidine) substrate (Mabtech) at RT and washing plates with distilled water immediately after spots became visible. Plates were dried at RT followed by imaging and spot enumeration using ImmunoSpot CTL analyzer.

### Single cell RNA sequencing and data analysis

MBCs were sorted from the lungs of allergen sensitized mice by FACS at day 21 or 77 (two biological replicates per timepoint), resuspended in PBS plus 0.05% bovine serum albumin (BSA, Thermo-Fisher Scientific), and counted using Trypan Blue and a hemacytometer. 2,500 cells per mouse were loaded into individual lanes for partitioning into Gel Bead-In-Emulsions (GEM) and barcoding RNA by 10x Genomics Chromium Controller (Loyola FACS Core). Reverse Transcription, cDNA amplification and library construction were done using Chromium Next GEM Single Cell kit 3’ v2 (10x Genomics). Library QC was done using Agilent 2100 Bioanalyzer and High Sensitivity DNA Kit (Loyola Genomics Core). Libraries were pooled and sequenced using paired-end sequencing (read 1 is 28 nt and read 2 is 90 nt in length) on two SP lanes with 28×90 cycles in a NovaSeq 6000 at the Roy J. Carver Biotechnology Center at University of Illinois at Urbana-Champaign. After demultiplexing, sequencing data were processed using CellRanger and aligned to GRCm38 reference genome (High Performance Computing in Biology, University of Illinois at Urbana-Champaign). Read count-UMI matrices were imported into R 4.2.2 and gene expression analysis was performed using the Seurat package v.4 (Satijalab.org) in R. QC was performed to filter out cells with high mitochondrial DNA counts (>8%), dying cells or poor-quality cells (gene count <500) or doublets (gene count >2500). Small numbers of contaminating plasmacytoid dendritic cells were removed based on cells expressing Klk1, Ramp1, Cldn5, or Cdh5. Gene expression values were normalized by total UMI counts per cell, multiplied by 10^4^ and log_10_ transformed. Highly variable genes were identified using the FindVariableFeature function in Seurat and linear transformed prior to dimensionality reduction by principal component analysis (PCA). Clustering and non-linear dimensionality reduction was performed using Uniform Manifold Approximation and Projection (UMAP) with 10 principal components to identify cell populations based on variable gene expression. Variable genes were exported as .csv files. Analysis of marker genes was done using FeaturePlot, VlnPlot, and DoHeatmap functions in Seurat. All four datasets were integrated for combined data analyses, or day 21 and 77 timepoints were integrated separately using IntegrateData and FindIntegrationAnchors functions in Seurat. Szymkiewics-Simpson similarity coefficient is calculated as the number of overlapping marker genes between two groups divided by the size of marker gene list with fewer number of genes^65^.

### Immunofluorescence microscopy

Perfused lungs were filled with a 50:50 mixture of PBS and Tissue-Tek optimal cutting temperature (OCT) compound via syringe and catheter in the trachea and embedded in OCT. OCT-embedded lungs were snap-frozen in liquid nitrogen-cooled methylbutane and stored at -80⁰ C. Frozen lungs were cut into 7-8 µm sections using a Cryostar NX50 cryostat (Thermo Fisher Scientific) and mounted on silane-treated glass slides (Sigma-Aldrich). Sections were rehydrated in PBS for 5 minutes at RT before fixing with 4% paraformaldehyde for 15 minutes at RT. Fixed sections were blocked in 10% rat serum (RS, Innovative Research, Inc.) in PBS for 1 hour at RT, then stained with antibodies against CD19 (1:500, APC, clone 6D5, BioLegend), CD4 (1:500, FITC, clone H129.19, BD), IgD (1:250, PE, clone 11-26c.2a, BioLegend) in PBS plus 10% RS overnight at 4⁰ C in a humidified chamber. Slides were washed in PBS, then FITC signal was amplified using anti-FITC HRP and AlexaFluor 488 Tyramide Signal Amplification Kit (Thermo Fisher Scientific). Stained slides were mounted with ProLong Gold Antifade Mountant with DAPI (Thermo Fisher Scientific). Z-stack images and panels were collected using the 20x objective on a DeltaVision wide field fluorescence microscope (Applied Precision, GE) with digital camera (CoolSNAP HQ, Photometrics) and analyzed using Imaris 8.4.1. Day 77 images were acquired using Zeiss 880 Airyscan with 10x 0.25NA objective for a tiled overview of the entire tissue, or a 63x 1.4NA objective for high-resolution imaging of peri-bronchiolar lymphoid aggregates. Raw data were processed using Airyscan processing in ‘auto strength’ mode. Linear adjustments for contrast and brightness were made to images using ImageJ.

### Parabiosis surgery

BALB/cByJ (CD45.2) and congenic CD45.1^+^ BALB/cByJ mice controlled for age and weight were surgically paired for 10 or 16 days following approved protocols by the Institutional Animal Care and Use Committee and guidelines of the Comparative Medicine Facility at Loyola University Chicago. Prior to surgery, mice were shaved, co-housed, and weighed. An incision was made in the skin of each mouse from front shoulder to rear leg under isoflurane inhalation anesthesia. Mice were joined at the elbow and knee with dissolvable sutures and the incision was closed with wound clips. Postoperative care included pain relief by slow-release buprenorphine, hydration by subcutaneous injection of 1 mL or intraperitoneal injection of 0.5 mL of 5% dextrose in saline, and addition of antibiotics to drinking water for 1 week to prevent infection (suspension of sulfamethoxazole and trimethoprim in water). In rechallenge experiments, mice were paired for 10 days, then pairs were rechallenged with nebulized OVA (1% in PBS) for 20 min/day for five consecutive days. Pairs were injected with anti-CD45 antibodies by tail vein injection 5 minutes prior to sacrifice to label intravascular lymphocytes, then tissues were harvested for analysis by flow cytometry.

### Figures and statistical analysis

Figures were prepared using Flowjo v10.4, GraphPad Prism v9.1.0, R v4.2.2, Adobe Illustrator 2023, and BioRender.com. Statistical analyses were performed with GraphPad Prism using one-way ANOVA with Tukey’s multiple comparisons test, unpaired two tailed Student’s *t*-test, or paired two tailed *t*-test with p values considered significant as indicated in figure legends. Data are presented as mean ± SD or mean ± SEM as indicated in figure legends. No data points were excluded.

## SUPPLEMENTAL INFORMATION

Supplemental information includes ten figures and one table on key resources.

## AUTHOR CONTRIBUTIONS

A.J.N. conceived the project, performed the experiments, generated figures and wrote the manuscript. B.K.T. and A.J.N. performed parabiosis experiments and contributed figures as well as comments to the manuscript. J.R.B. performed imaging analyses on day 77 sensitized mouse lungs and contributed figures as well as comments to the manuscript. D.K.S. supervised work and contributed comments to the manuscript. Y.L.W. conceived the project, supervised work, and wrote the manuscript.

## ACKNOWLEDGEMENTS

We are grateful to Patricia Simms and Robert Ladd at the Loyola FACS core facility for their technical support on cell sorting. We thank Dr. Adarsh Dharan and Dr. Edward Campbell for their technical support with immunofluorescence microscopy. We thank staff at the Loyola Comparative Medicine Facility for their excellent work in animal husbandry. We thank members of the Wu Lab for their critical comments and helpful support. This work was supported by grants from the National Institutes of Health R01HL165120 and R21AI159456 to Y.L.W., and A.J.N. was supported by F31HL156459 and T32AI007508.

