## Supplemental Materials for "Lung-resident memory B cells maintain allergic IgE responses in the respiratory tract"

#### SUPPLEMENTAL INFORMATION

Supplemental information includes ten figures and one table on key resources.

#### SUPPLEMENTAL FIGURE LEGENDS

**Figure S1. Allergen inhalation induces eosinophilia, IgE and IgG1.**

**(A)** Numbers of eosinophils in the lungs on day 21 by flow cytometry analyses.

**(B)** Measurements of OVA-specific IgE (left) and IgG1 (right) in sera on day 21 by ELISA.

Data are represented as mean ± SD. Student's *t*-test; *, $p<0.05$.

**Figure S2. Germinal center responses to allergen inhalation.**

**(A)** Representative plots and frequencies of germinal center (GC) B cells on day 21 by flow cytometry analyses.

**(B)** Representative plots and frequencies of T follicular helper ($T_{FH}$) cells on day 21 by flow cytometry analyses.

Data are represented as mean ± SD. Representative data from three independent experiments.

**Figure S3. Effects of FTY720 on IgE-switching B cells in the lungs.**

**(A)** Intravascular labeling of lung B cells with anti-CD45 antibodies to identify B cells in the lung tissue and the lung vasculature on day 21 after 3 days of treatment with saline (top) or FTY720 (bottom). Gated on live $B220^+$ cells. Dashed lines indicate gating used to generate percentages.

**(B)** Representative flow cytometry plots showing frequencies of Iε-tdTomato$^+$ B cells (live i.v. CD45$^-$ B220$^+$) in the lung tissue of saline (left) or FTY720-treated (right) mice on day 21.

**(C)** Numbers of Iε-tdTomato$^+$ B cells in the lung tissue (left) or the lung vasculature (right) of saline or FTY720-treated mice on day 21. Data are represented as mean ± SD, Student's *t*-test.

**Figure S4. Identification of IL-4-producing T cells.**

**(A)** Frequencies of T follicular helper ($T_{FH}$) cells among CD4$^+$ effector T cells in each tissue from sensitized 4Get mice on day 21 by flow cytometry analyses (top). Frequencies of GFP$^+$ $T_{FH}$ in the same mice (bottom).

**(B)** Frequencies of T helper type 2 ($T_H2$) cells among total CD4$^+$ T cells in each tissue on day 21 by flow cytometry.

**(C)** Frequencies of $T_H2$ cells among GFP$^+$ CD4$^+$ T cells in each tissue on day 21 by flow cytometry.

Representative data from two independent experiments. Data are represented as mean ± SD, one-way ANOVA with Tukey's multiple comparison's test; ns, p>0.05; *, p<0.05; ** p<0.01; ****, p<0.0001.

**Figure S5. Requirement for antigen and T cells for IgE class switching in coculture experiments.**

Frequency of Iε-tdTomato$^+$ MBCs after 3 days in culture without (left) or with CD4$^+$ T cells from the lungs (middle) or spleen (right) in the presence (top) or absence (bottom) of OVA.

**Figure S6. Effects of FTY720 or anti-CD20 depletion on the numbers of B cells in the lung tissue or vasculature.**

**(A)** Numbers of naïve B cells, $IgM^+$ MBCs, or $IgG1^+$ MBCs in the lung tissue or vasculature after 8 days of treatment with saline or FTY720.

**(B)** Numbers of naïve B cells, $IgM^+$ MBCs, or $IgG1^+$ MBCs in the lung tissue or vasculature 8 days after the injection of 50 μg of isotype control (IgG2a) or anti-CD20 antibodies.

Data are representative of two independent experiments with 8 mice, and presented as mean ± SD, one-way ANOVA with Tukey's multiple comparison's test.

**Figure S7. Lung eosinophils after allergen sensitization.**

Representative plots showing frequencies of eosinophils in the lungs (left) and numbers of lung eosinophils (right) at each timepoint.

**Figure S8. Heatmaps of marker gene expression in day 21 and day 77 single cell RNA-seq libraries.**

**(A)** Heatmap showing scaled expression of the top 10 marker genes from each cluster in the day 21 libraries.

**(B)** Heatmap showing scaled expression of the top 10 marker genes from each cluster in the day 77 libraries.

Each time point contains two individual libraries.

**Figure S9. Expression of day 77 cluster marker genes in the day 21 libraries.**

Violin plots showing scaled expression of marker genes from day 77 Cluster 0 (top), Cluster 1 (middle) or Cluster 2 (bottom) by Clusters 0-3 in the day 21 libraries.

**Figure S10. OVA-specific IgG1 antibody-secreting cells after rechallenge with OVA.**

Detection of OVA-specific IgG1 antibody-secreting cells (ASCs) after OVA rechallenge of PBS control (left) or RWP+OVA-sensitized (right) mice by ELISpot. Duplicate wells for each tissue are shown. Representative of two independent experiments with 4 or 8 mice.

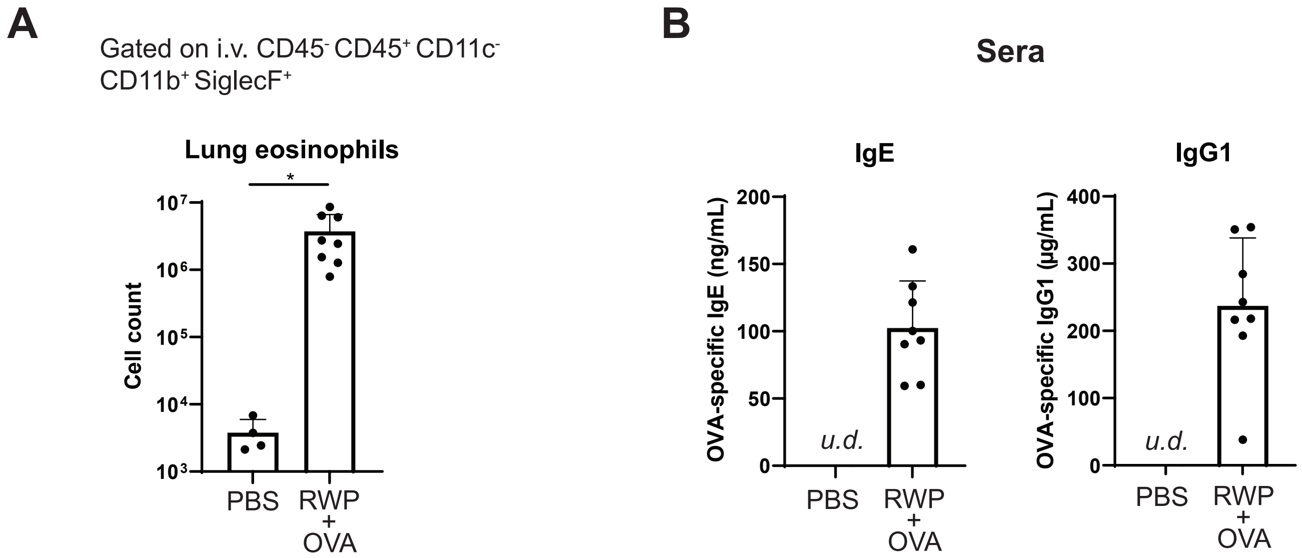


**Nelson et al. Figure S1**

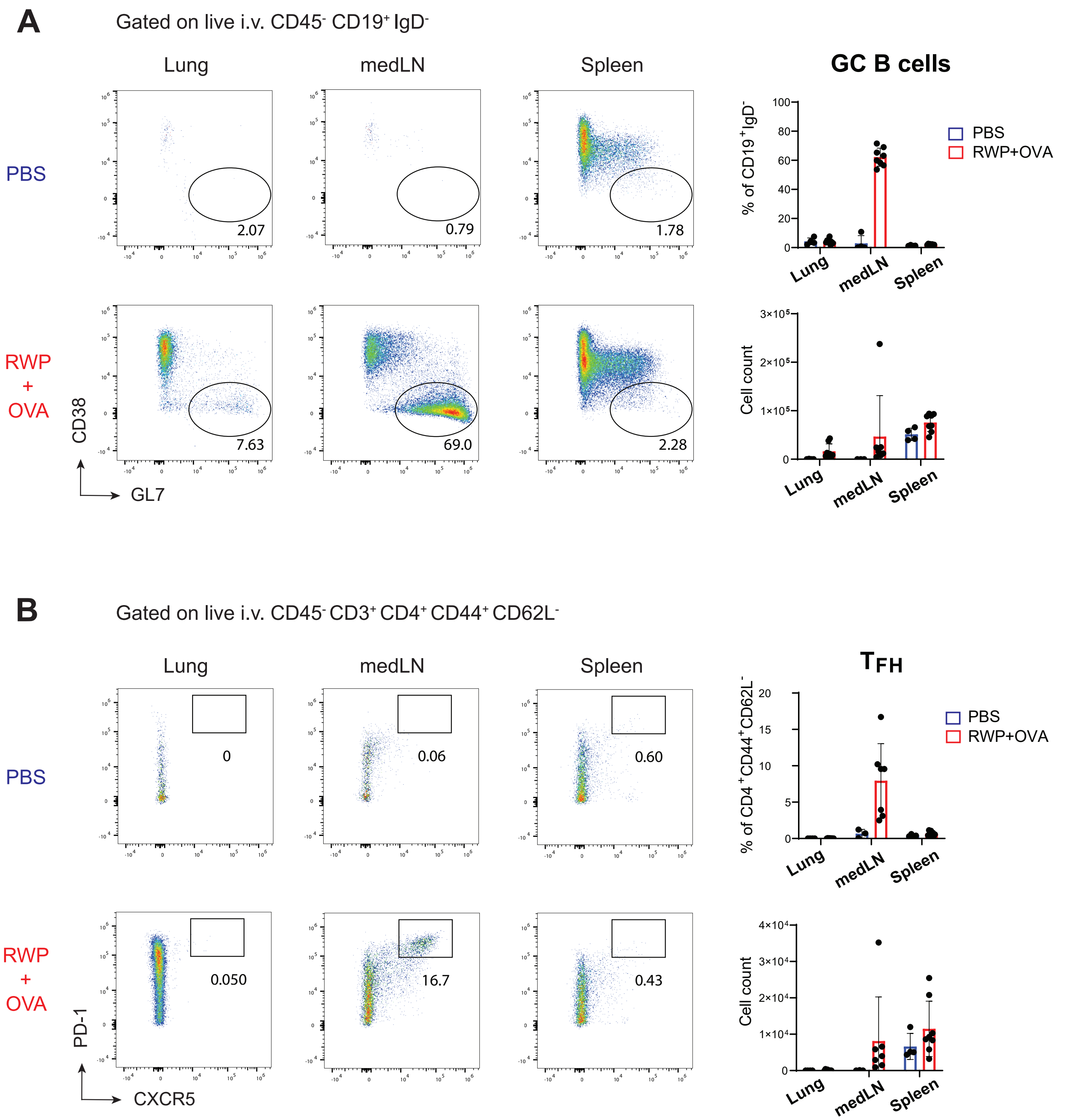


Nelson et al. Figure S2

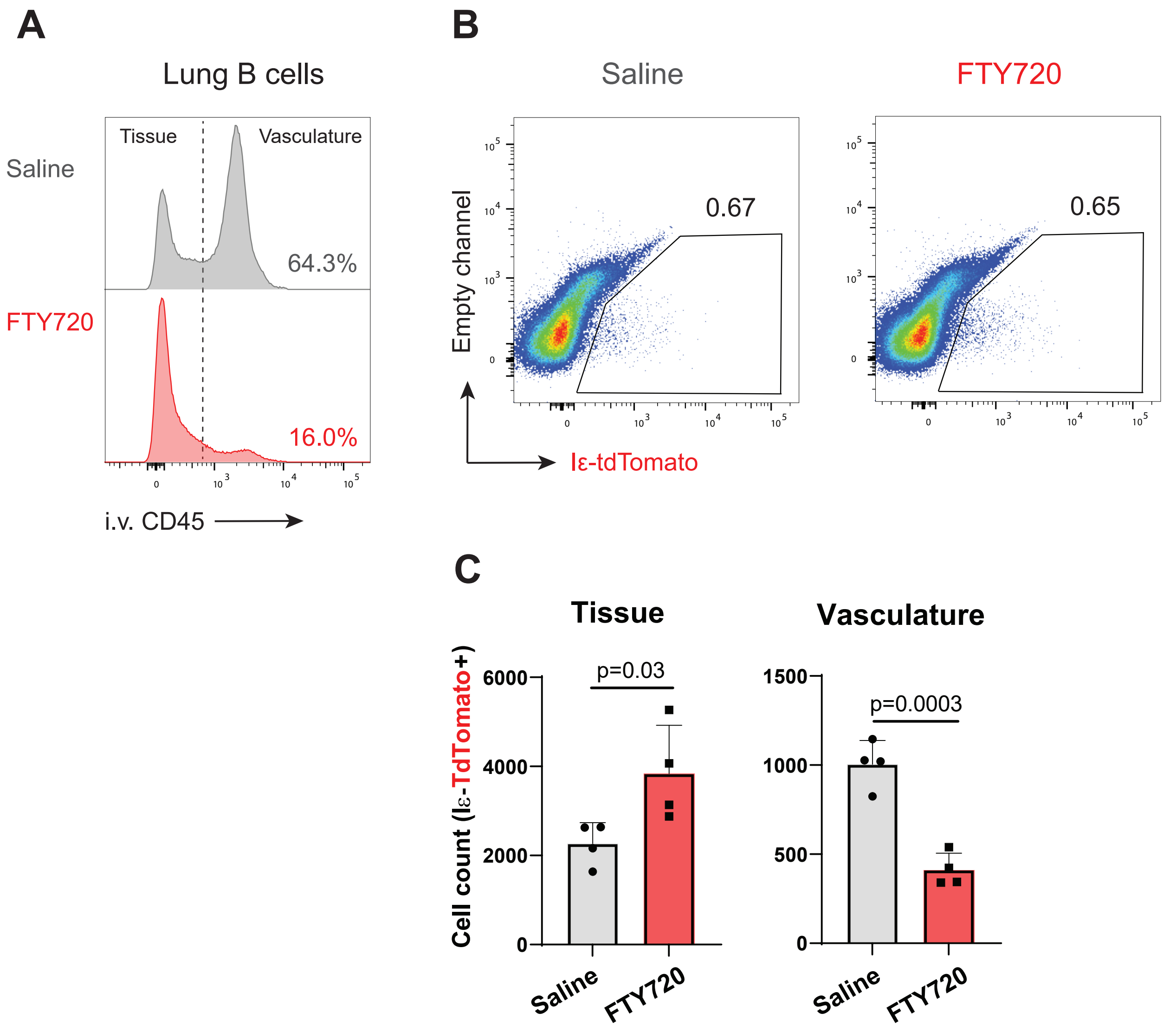


Nelson et al. Figure S3

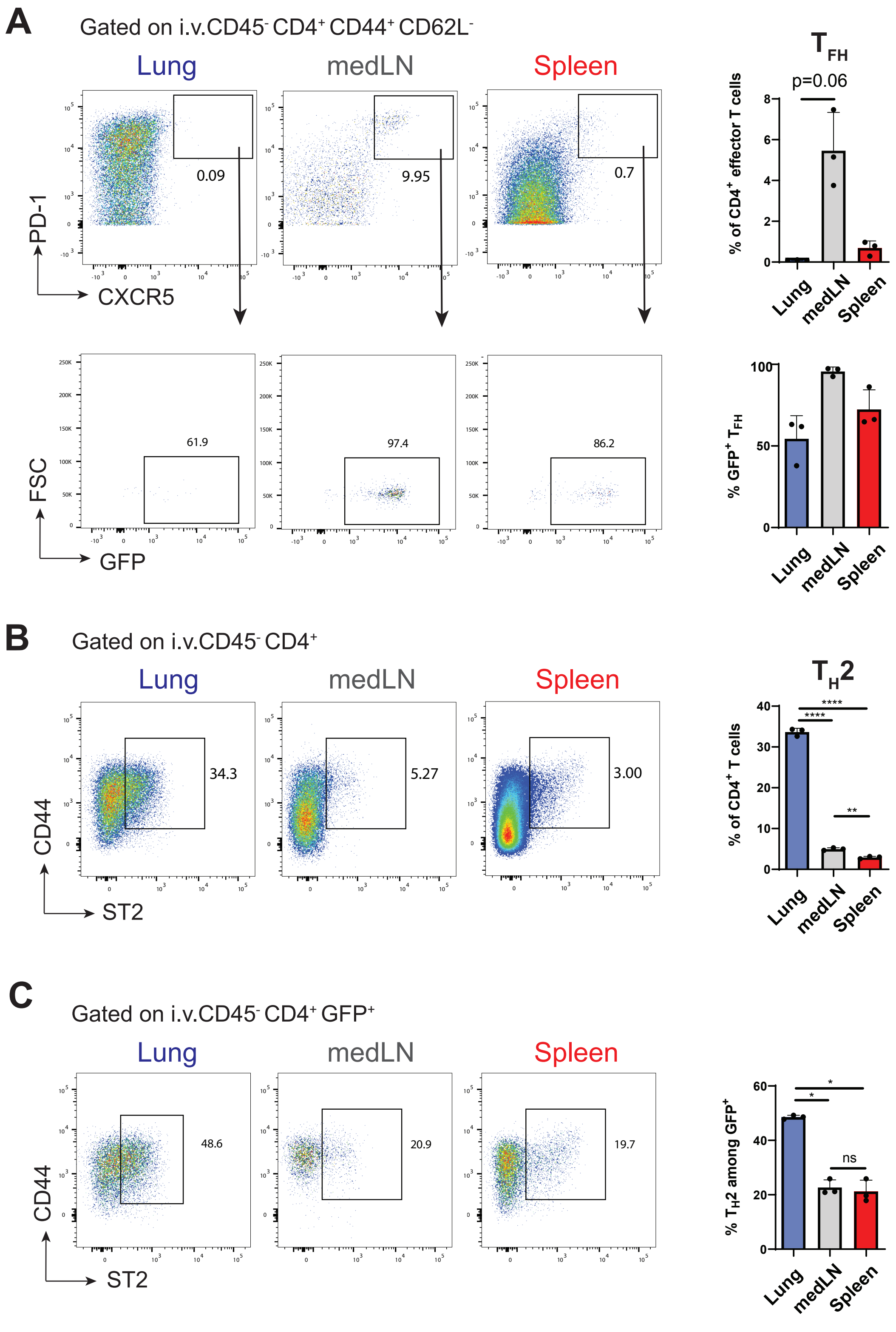


Nelson et al. Figure S4

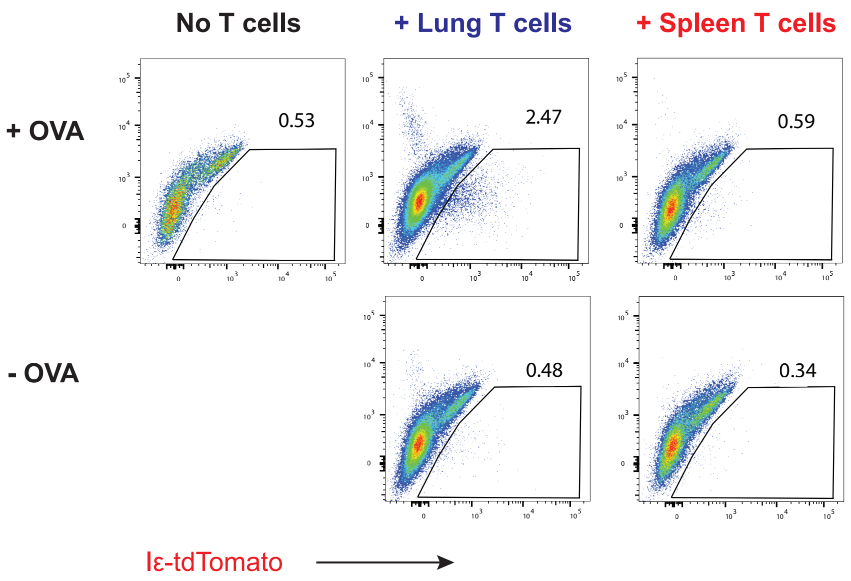


**Nelson et al. Figure S5**

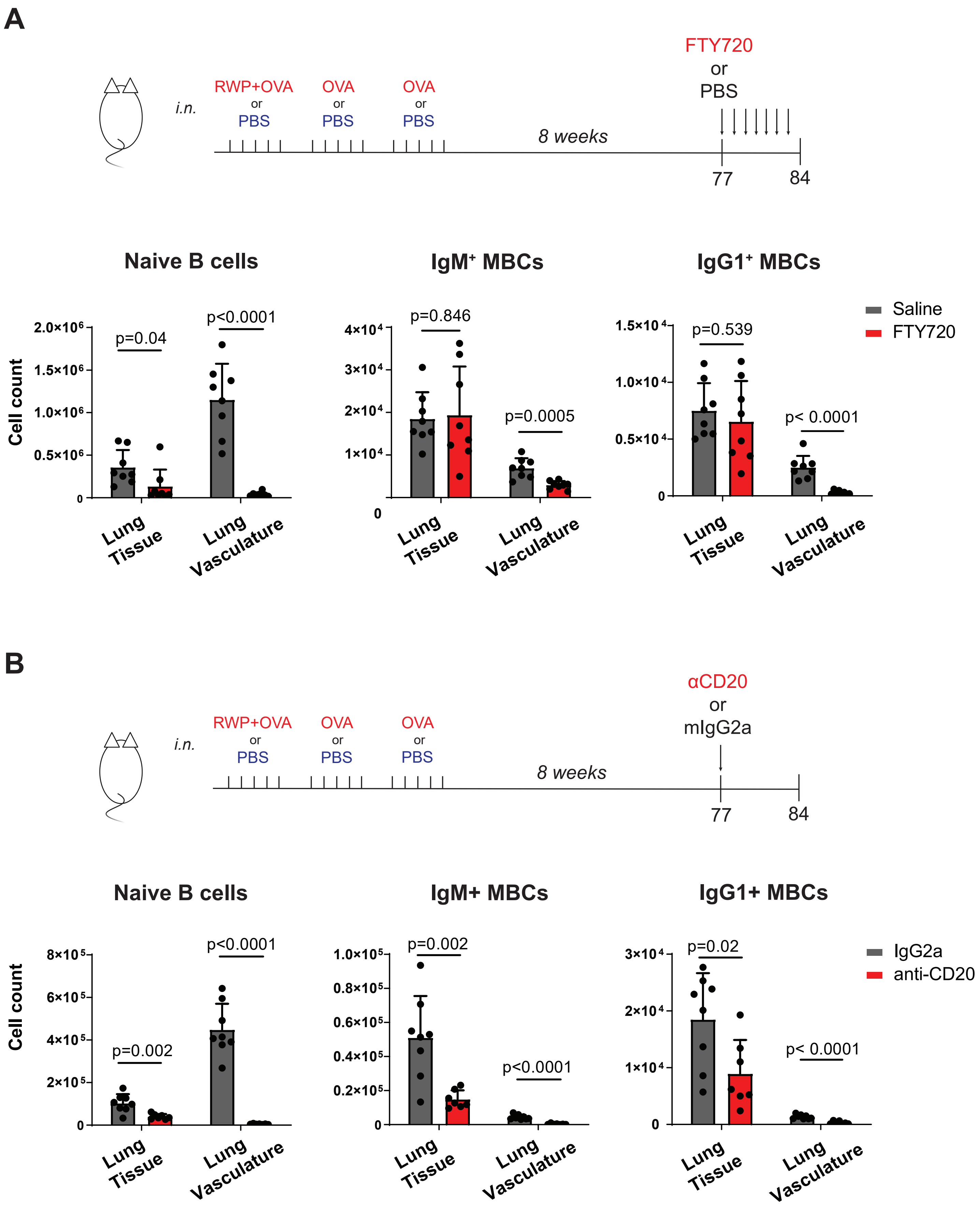


**Nelson et al. Figure S6**

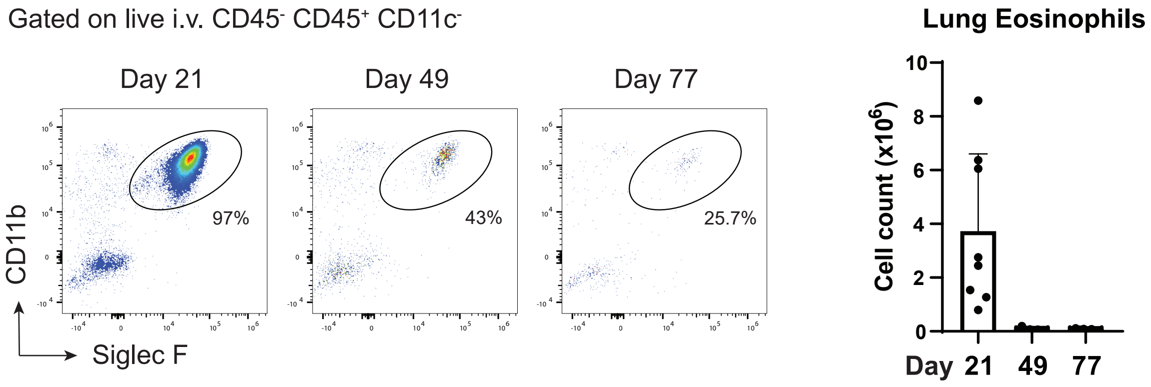


Nelson et al. Figure S7

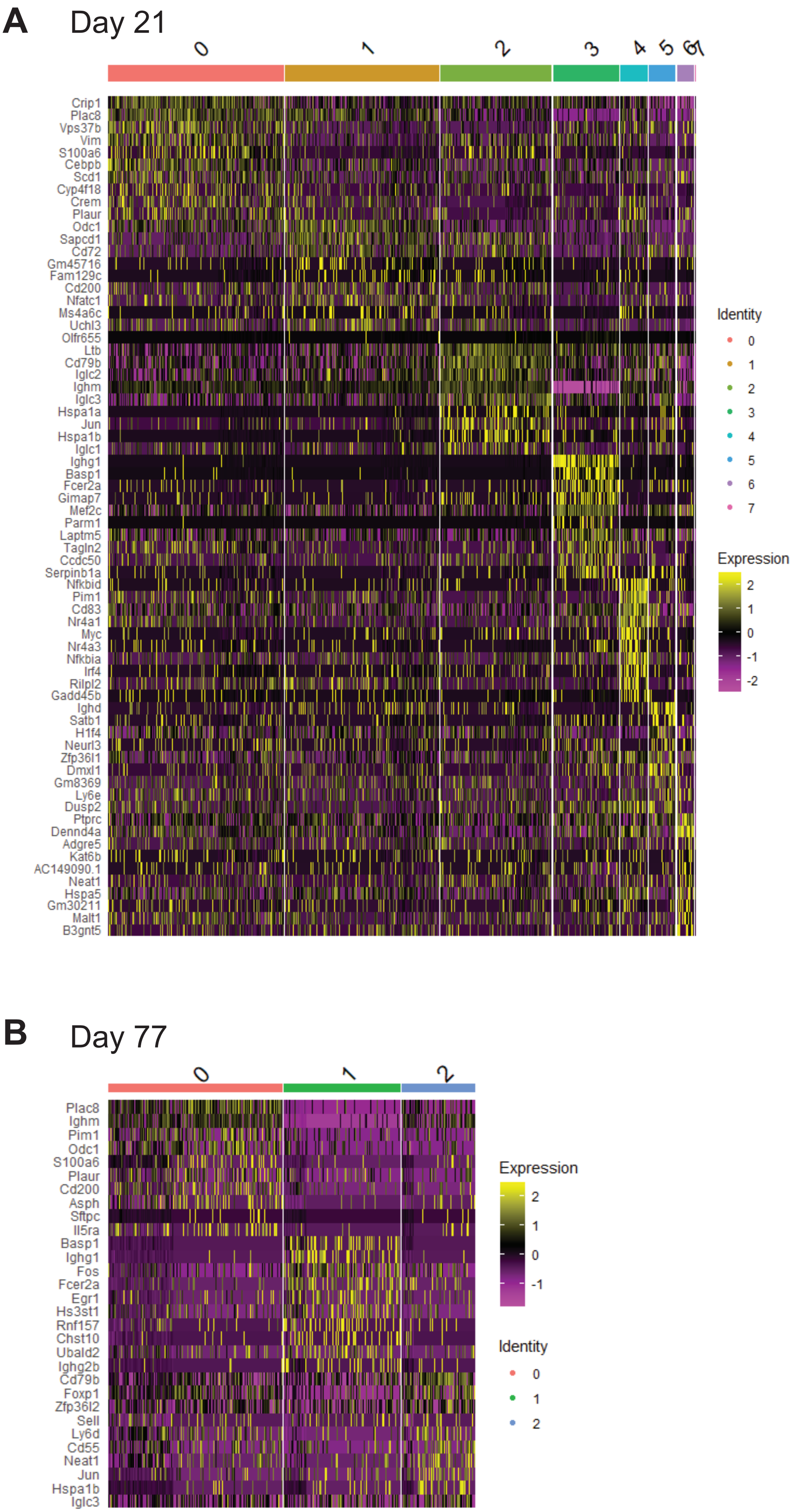


**Nelson et al. Figure S8**

### Day 21 Libraries

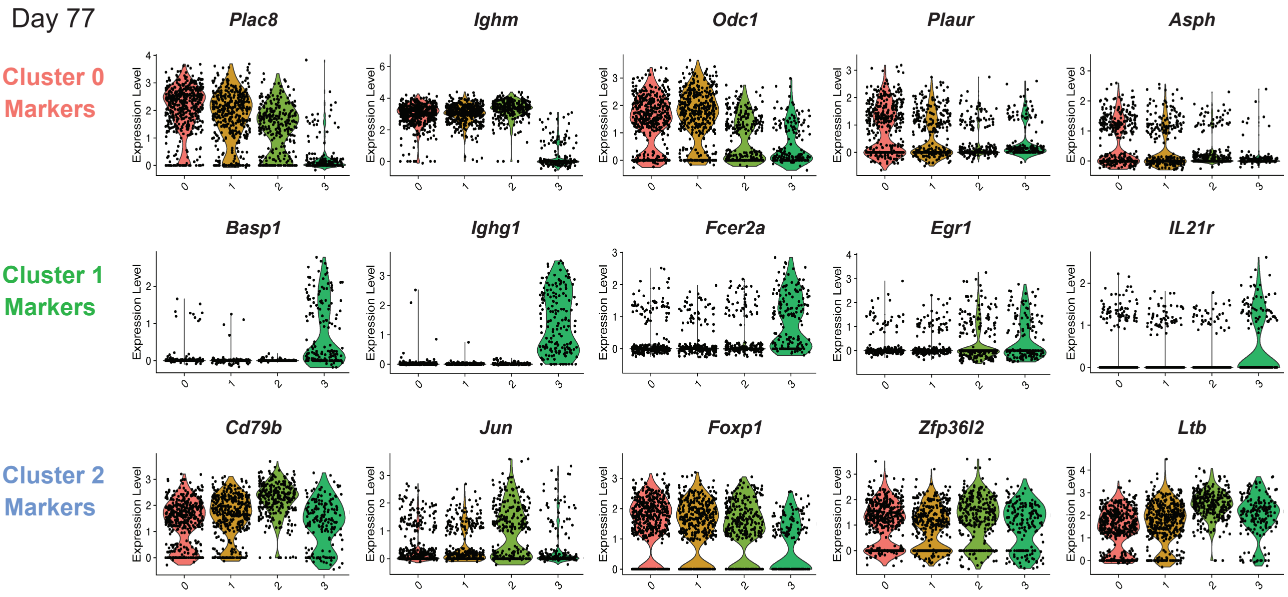


**Nelson et al. Figure S9**

#### OVA-specific IgG1 ASCs

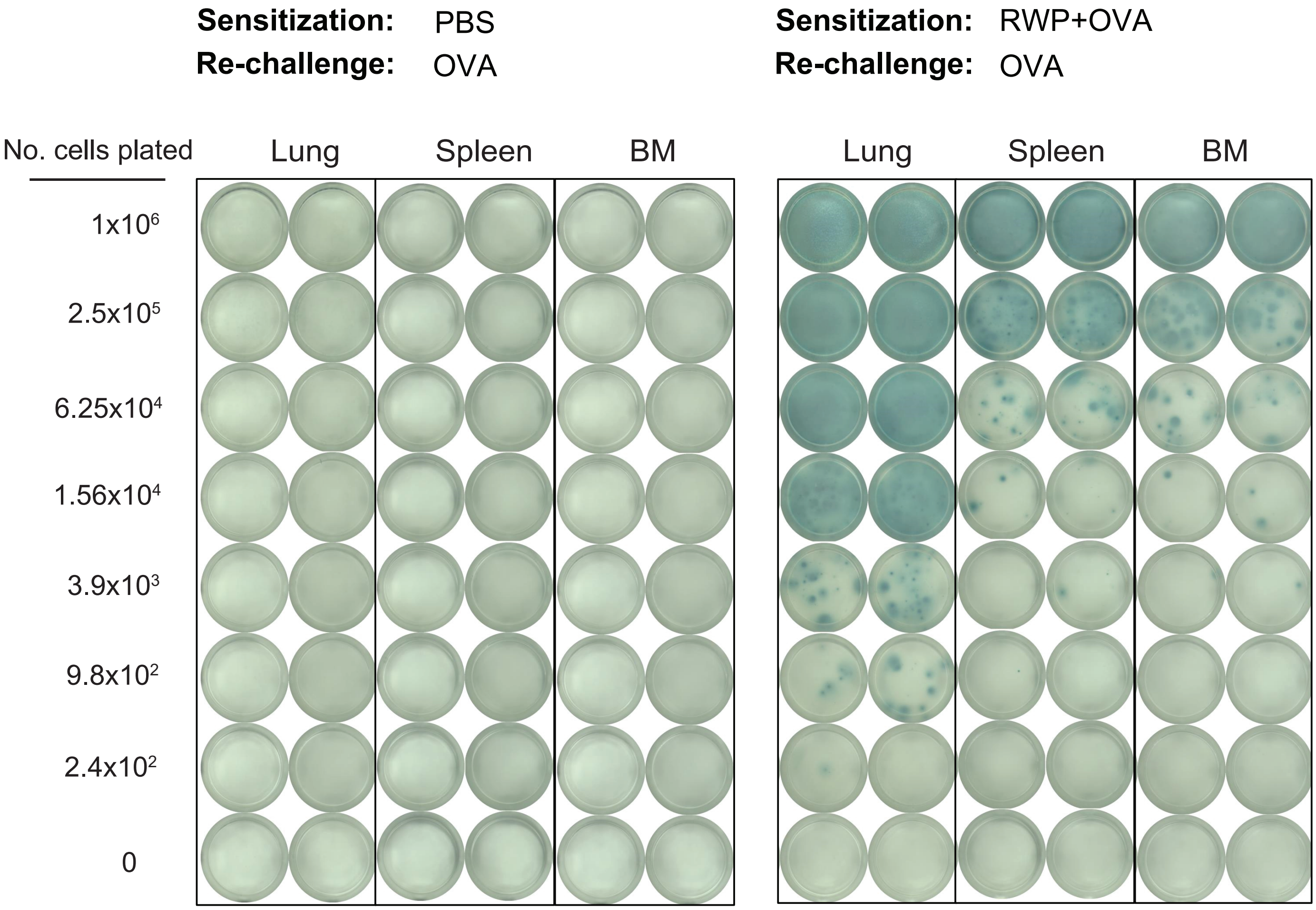


**Nelson et al. Figure S10**

**Table S1. List of antibodies used in this study.**

| Target | Format | Clone | Source | Catalog number | Dilution |
|---|---|---|---|---|---|
| CD19 | Spark Blue 550 | 6D5 | BioLegend | 115566 | 1:800 |
| CD19 | APC | 6D5 | BioLegend | 115511 | 1:800 |
| B220 | PacBlue | RA3-6B2 | BioLegend | 103230 | 1:400 |
| CD38 | FITC | 90 | BioLegend | 102705 | 1:800 |
| CD38 | APC-cy7 | 90 | BioLegend | 102727 | 1:800 |
| IgD | BUV395 | 11-26C.1 | BD | 564274 | 1:400 |
| IgD | BV650 | 11-26C.1 | BioLegend | 405721 | 1:400 |
| IgD | PE | 11-26C.1 | BioLegend | 405705 | 1:400 |
| GL7 | PacBlue | GL7 | BioLegend | 144614 | 1:200 |
| GL7 | PE | GL7 | BioLegend | 144608 | 1:200 |
| IgM | PercP-cy5.5 | RMM-1 | BioLegend | 406512 | 1:50 |
| IgG1 | APC-cy7 | RMG1-1 | BioLegend | 406619 | 1:1000 |
| IgG1 | APC | RMG1-1 | BioLegend | 406609 | 1:1000 |
| CD80 | BV605 | 16-10A1 | BioLegend | 104729 | 1:200 |
| CD80 | PE-cy7 | 16-10A1 | BioLegend | 10473 | 1:200 |
| PD-L2 | BUV563 | TY25 | BD | 741431 | 1:1000 |
| PD-L2 | APC | TY25 | BioLegend | 107210 | 1:1000 |
| CCR6 | BV785 | 29-2L17 | BioLegend | 129823 | 1:100 |
| CD23 | BUV737 | B3B4 | BD | 749668 | 1:400 |
| CCR6 | BV785 | 29-2L17 | BioLegend | 129823 | 1:100 |
| CD23 | BUV737 | B3B4 | BD | 749668 | 1:400 |
| CD73 | BV421 | TY/11.8 | BioLegend | 127217 | 1:200 |
| IL-21Ra | PE-cy5 | 4A9 | BioLegend | 131908 | 1:50 |
| IL-5R | BUV805 | T21 | BD | 742063 | 1:200 |
| CD69 | BV711 | H1.2F3 | BioLegend | 104537 | 1:200 |
| CD29 | Alexa 700 | HMβ1-1 | BioLegend | 102218 | 1:400 |
| CD11a | BUV661 | 2D7 | BD | 741469 | 1:200 |
| CD49d | PE/Dazzle 594 | R1-2 | BioLegend | 103626 | 1:100 |
| CD3 | APC | 145-2C11 | BioLegend | 100312 | 1:200 |
| CD4 | FITC | H129.19 | BD | 553650 | 1:1000 |
| CD45 | BUV615 | I3/2.3 | BD | 752418 | 1:400 |
| CD44 | BV785 | IM7 | BioLegend | 103041 | 1:200 |
| CD62L | PacBlue | MEL-14 | BioLegend | 104424 | 1:1000 |
| CD62L | Biotin | MEL-14 | BioLegend | 104403 | |
| ST2 | PerCP-eFlour710 | RMST2-2 | Invitrogen | 46-9335-80 | 1:200 |
| PD-1 | PE | 29F.1A12 | BioLegend | 135206 | 1:500 |
| CD11b | Spark YG 593 | M1/70 | BioLegend | 101281 | 1:800 |
| CD11b | APC-cy7 | M1/70 | BioLegend | 101225 | 1:800 |

| | | | | | |
|---|---|---|---|---|---|
| CD11c | Spark Blue 550 | N418 | BioLegend | 117365 | 1:200 |
| CD11c | PerCP | N418 | BioLegend | 117325 | 1:200 |
| Siglec F | BV480 | E50-2440 | BD | 552125 | 1:400 |
| Siglec F | PE | S17007L | BioLegend | 155505 | 1:400 |
| CXCR5 | Biotin | L138D7 | BioLegend | 145510 | 1:200 |
| CD45 | PacBlue | 30-F11 | BioLegend | 103125 | |
| CD45 | BV510 | 30-F11 | BioLegend | 103137 | |
